# Optimizing phenotype scale improves genetic analyses in large-scale biobanks

**DOI:** 10.64898/2026.05.04.722531

**Authors:** Zhenhong Huang, Manuela Costantino, Andy Dahl

## Abstract

Large-scale biobanks have enabled increasingly complicated genetic analyses across thousands of phenotypes. However, studies rarely consider the appropriate phenotype measurement scale, a problem that can drastically affect inferences on genetic architecture. Here, we introduce SIQReg, a practical solution to this classical problem, which learns a data-driven phenotype scale by minimizing heterogeneity across phenotype quantiles. Applied to complex traits in UK Biobank, SIQReg rejects the default scale for 24/25 traits. Generally, SIQReg scales lie between default and logarithmic, indicating that default-scale traits are neither purely additive nor purely multiplicative. We show that SIQReg improves both non-additive and additive genetic analyses. SIQReg eliminates most non-additive genetic signals (such as 97% of vQTL and 76% of quantile-dependent TWAS genes), indicating they may be statistical artifacts, while preserving biologically plausible non-additive signals. Simultaneously, SIQReg improves power to detect additive signals, increasing GWAS loci, TWAS genes, and PGS prediction accuracy by 11%, 13%, and 10%, respectively, and identifies 50% more high-risk individuals. These gains replicate across ancestry groups. Our results establish SIQReg as a principled approach to phenotype scale transformation that improves genetic analyses of complex traits.

## Introduction

Large-scale biobanks have reshaped human genetics by enabling genome-wide association studies (GWAS) with unprecedented numbers of individuals and phenotypes^1,2^. These studies have been remarkably successful in identifying loci associated with complex traits using additive models, as well as suggesting causal genes^3,4^ and training polygenic scores (PGS) to predict disease risk^5,6^ and identify high-risk individuals^7^. While these systematic scans across traits have provided invaluable resources, their broad scope limits opportunities to optimize analyses for individual phenotypes^8,9^.

One popular direction to expand beyond standard GWAS is to model non-additive genetic effects. Non-additive models have become increasingly common due to the proliferation of large-scale biobank-based datasets, which provide the power needed to detect non-additive genetic effects. For example, PGSxE tests of gene-environment interaction (GxE) are increasingly common due to their simplicity and power^10–14^, and variance quantitative trait loci (vQTL) have been widely reported as a signal of underlying genetic interactions^15,16^. More recently, studies using quantile regression have identified genetic effects that differ across quantiles of the phenotype distribution^17–24^.

However, growing evidence suggests that non-additive genetic tests can depend heavily on the phenotype scale. When an additive trait is measured on a non-additive scale, nearly all effects will appear non-additive with sufficient power. Such scale-induced heterogeneity is arguably a statistical artifact and obfuscates the search for scale-independent heterogeneity^25–27^. In practice, domain-specific studies often carefully consider the appropriate scale, such as logarithmic, rank-based inverse normal transformation (RINT)^28–32^, or more tailored transformations^33^. In biobank-based multi-trait analyses, however, phenotype scale is rarely considered or stress-tested. Although this issue of phenotype scale has been discussed for nearly a century^34–36^, there is still no general solution.

Here, we address this gap with SIQReg (scale-independent quantile regression), which learns a phenotype scale to minimize scale-dependent heterogeneity. The key insight underlying SIQReg is that, on an appropriate scale, additive genetic effect sizes are identical across the phenotype distribution. SIQReg therefore seeks a scale that minimizes this variation. We apply SIQReg to 25 phenotypes in UK Biobank and reject the default scale for 24/25. We find that SIQReg removes substantial genetic heterogeneity while simultaneously improving power for additive genetic effects. Our findings show that SIQReg is a simple and general approach to improve genetic analyses.

## Results

### Overview of SIQReg

When a phenotype is measured on a non-additive scale, genetic effects that are truly constant will appear to vary across the phenotype distribution, creating non-additive genetic signals that are statistically significant but biologically meaningless. Specifically, suppose that there exists a purely additive latent phenotype, *y*\*, but we are provided the default-scale phenotype, *y* (**Fig. 1**). We assume that 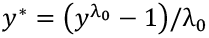 is a Box-Cox transformation of *y* with exponent *λ*_0_, which we call the latent scale transformation parameter; importantly, this simplifies to the default scale when *λ*_0_ = 1, and *λ*_0_ = 0 corresponds to the log scale. In general, *λ*_0_ ≠ 1 implies that the effect of additive genetic variant *G* will appear quantile-dependent on the default scale: 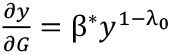 (**Note S1.1**). Quantile-dependent effects thus provide a natural test of phenotype scale validity. More generally, we show that quantile regression is a flexible and agnostic test of genetic heterogeneity that captures signals including GxE and vQTL (**Note S1.3)**.

**Figure 1.**
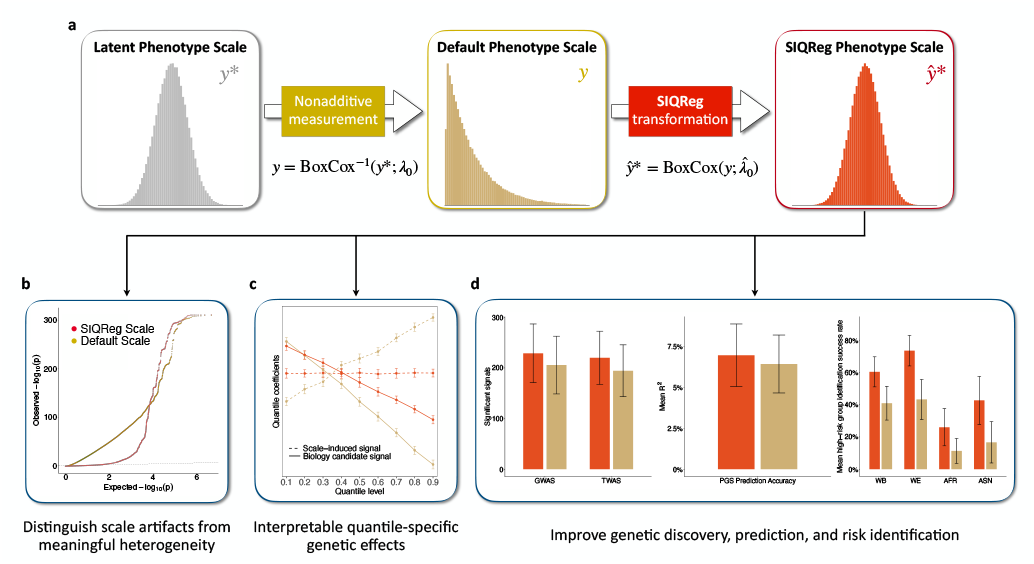
SIQReg recovers latent additive phenotype scale to improve genetic analyses. (a) The latent additive phenotype, *y*\*, is observed on some non-additive default phenotype scale, *y*, which induces non-additive effects. SIQReg estimates a transformed phenotype scale, 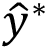, that more closely approximates the latent scale. In practice, SIQReg optimizes over Box-Cox transformations. (b) SIQReg removes scale-dependent genetic heterogeneity signals (such as vQTL and quantile-dependent genetic effects) while preserving signals that are more likely to reflect biology. (c) SIQReg distinguishes scale-induced heterogeneity from scale-independent quantile-specific genetic effects. (d) SIQReg improves power for additive genetic analyses, including genetic discovery, polygenic prediction accuracy, and identification of high-risk individuals.

Mathematically, SIQReg identifies the scale transformation 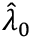 by minimizing quantile-dependent effect heterogeneity (**Methods, Note S1.2**):

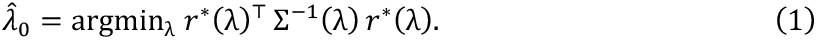

where *r*\*(*λ*) is the residual vector of quantile regression coefficients across quantile levels and Σ(*λ*) is its covariance matrix. In practice, this procedure is applied to multiple covariates, and the resulting 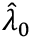 are meta-analyzed.

The value of 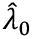 is informative in itself: deviations from 1 indicate that default-scale analyses will produce scale-dependent artifacts, and values near 0 suggest a simple log-transformation, i.e., that default-scale genetic effects are multiplicative. But the primary utility of SIQReg is to improve downstream genetic analyses by using 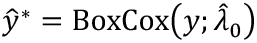 in place of *y*: by design, SIQReg removes scale-dependent heterogeneity, but we also prove that it improves power to detect additive effects (**Note S1.4**).

### SIQReg recovers the latent scale and improves genetic analyses in simulations

We conducted simulations to test the ability of SIQReg to recover the correct latent scale. As expected, when we simulated default-scale phenotypes that had been Box-Cox transformed with parameter *λ*_0_, SIQReg unbiasedly estimated this parameter, with increasing accuracy as sample size grows (**Fig. 2a**). Because SIQReg estimates *λ*_0_ by minimizing quantile-dependent covariate effects, estimation accuracy also improved with the covariate effect size (**Fig. S1**).

**Figure 2.**
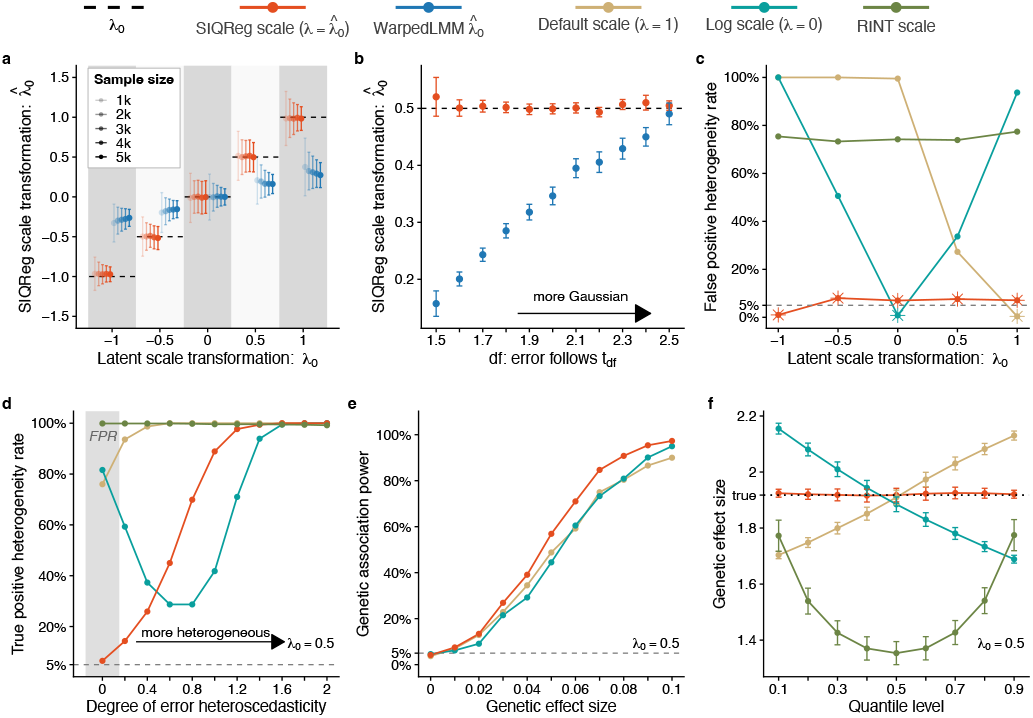
Simulations demonstrate that SIQReg recovers latent additive scales and improves downstream genetic analyses. (a) Impact of sample size on SIQReg and WarpedLMM scale estimates under t-distributed errors, evaluated across five values of latent scale parameters, *λ*_0_. (b) Impact of non-Gaussianity on SIQReg and WarpedLMM estimates, parameterized by varying t-distribution degrees of freedom. (c) False-positive rates for testing quantile-dependent heterogeneity; * indicates the values of *λ*_0_ where an assumed scale holds. (d) True-positive rate to detect quantile-dependent heterogeneity compared to the strength of quantile-dependence, parameterized by heteroscedasticity; the grey region indicates the null (no true heterogeneity), showing that default, log, and RINT are severely inflated. (e) SIQReg improves power to detect additive genetic effects. (f) Illustrative example of quantile-dependent genetic effects across scales, showing that the existence and qualitative nature of heterogeneity vary substantially across scales. Error bars represent the standard error of the mean across 1,000 simulation replicates in (a) and (b); error bars in (f) represent the 95% confidence intervals for the point estimates.

We compared SIQReg to the likelihood-based approach assuming Gaussian noise, similar to WarpedLMM^33^ (**Methods**). Both WarpedLMM and SIQReg perform well when the latent scale has Gaussian errors. However, only SIQReg remained unbiased in the presence of non-Gaussian errors (**Fig. 2b**). Intuitively, WarpedLMM seeks the scale with approximately Gaussian residuals, which fails in the realistic scenario where the additive scale has over-dispersed errors.

We next asked how SIQReg impacted downstream quantile regression tests of genetic heterogeneity under an additive model. SIQReg gave calibrated false-positive rates across latent scales, while the default and log scales were miscalibrated except under their assumed models (*λ*_0_ = 0 and *λ*_0_ = 1, respectively, **Fig. 2c**). We found that RINT was severely inflated regardless of the value of *λ*_0_; by contrast, RINT is only mildly inflated for realistic GxE^26^, illustrating how quantile regression is more informative of phenotype scale. We then tested power to detect scale-independent heterogeneity by simulating quantile-dependent effects on the latent scale, which confirmed that SIQReg preserves power to detect true scale-independent heterogeneity (**Fig. 2d**). We also found that SIQReg improved power to detect additive effects when *λ*_0_ ≠ 1 **(Fig. 2e**), confirming our theoretical results (**Note S1.4**).

Finally, we illustrated how phenotype scale can severely bias quantile regression results in the case of an additive genetic effect (**Fig. 2f)**. As desired, the genetic effect is constant across phenotype quantiles on the latent scale and the SIQReg scale. However, the effect increases on the log scale; decreases on the original scale; and is U-shaped on the RINT scale. In this example, the biological conclusion is entirely determined by the choice of measurement scale.

### SIQReg rejects default phenotype scales in UK Biobank

We applied SIQReg to estimate *λ*_0_ across 25 phenotypes in unrelated White British individuals in UK Biobank (**Methods**). We used 43 covariates as SIQReg inputs to learn *λ*_0_, including sex, age, age2, and genetic principal components. We found that 24/25 SIQReg scale estimates, 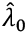, were significantly below 1 (p < 0.05, **Fig. 3a**), clearly demonstrating that the default scale provided by UK Biobank is usually suboptimal. On average across traits, we found 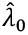 = 0.104 (*SE* = 0.055), suggesting the log scale generally outperforms the default scale; nonetheless, 7/25 SIQReg estimates lie strictly between the log and default scales. While a simple interpretation is that default-scale phenotypes are square-root additive (i.e., *λ*_0_ = 0.5), we propose a more mechanistic interpretation in which each complex trait reflects a mix of additive and multiplicative pathways (**Note S1.5**).

**Figure 3.**
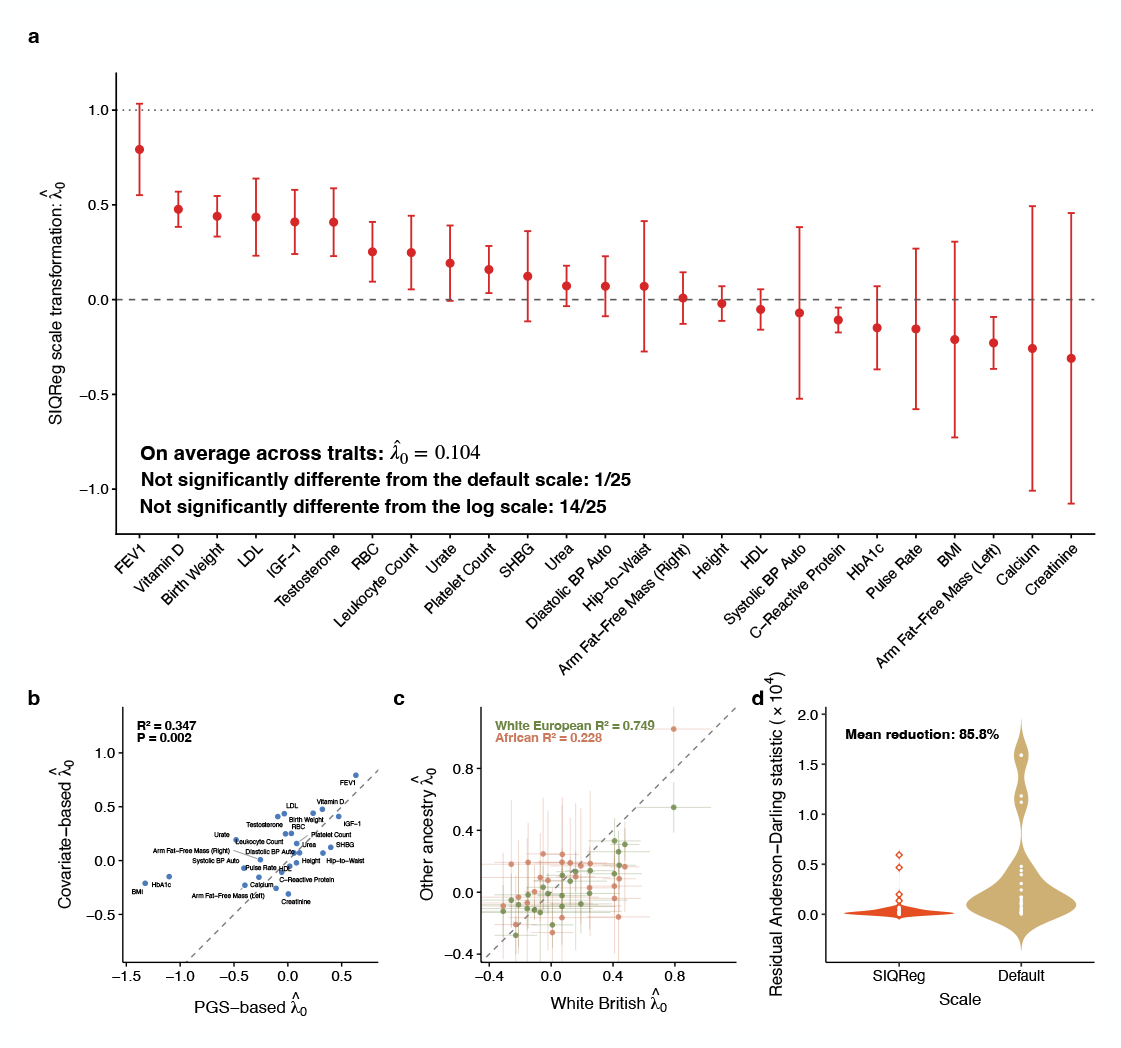
SIQReg robustly rejects the default phenotype scale in UK Biobank. (a) SIQReg estimates of the latent scale parameter 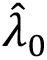 across 25 complex traits (**Table S1**). (b) 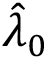 using PGS as input (x-axis) are concordant with primary SIQReg estimates of 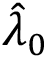, which use 43 covariates as input (y-axis). (c) 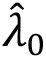 is consistent across ancestry groups in UK Biobank. (d) SIQReg substantially reduces residual Anderson–Darling statistics relative to the default scale; for clarity, extreme default-scale Anderson–Darling statistics were capped for C-reactive protein and HbA1c. Error bars represent the 95% confidence intervals and are truncated in (b) for clarity.

We validated these estimates of *λ*_0_ with several secondary analyses. First, we estimated *λ*_0_ using PGS as input to SIQReg to learn *λ*_0_ instead of covariates. Despite the weaker effect sizes of PGS, the estimated 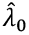 were generally concordant (*R*^2^ = 0.347, *p* = 0.002, **Fig. 3b**). Second, we estimated 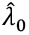 in White European, African, and Asian ancestry groups in UK Biobank and found highly consistent estimates with White British (*R*^2^ = 0.749, 0.228, 0.324, respectively, **Fig. 3c, Fig. S2**); the lower *R*^2^ in African and Asian groups is likely due to their smaller sample sizes. Third, we confirmed that our estimates are robust to outliers by excluding individuals with phenotypes more than 6 standard deviations from the mean^37^ (*R*^2^ = 0.908, **Fig. S3**). Fourth, *λ*_0_ estimates from SIQReg and WarpedLMM were also correlated (*R*^2^ = 0.634, **Fig. S4**), though simulations showed that SIQReg estimates are more robust. Finally, we tested robustness to the number of input covariates and found that 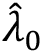 was stable so long as we included at least 13 of the 43 total covariates (**Fig. S5-6**). Together, these results show that SIQReg scale estimates are robust.

While SIQReg scales phenotypes to minimize scale-dependent heterogeneity, we asked how SIQReg impacted two other classical criteria for phenotype scale: non-Gaussianity and heteroscedasticity^35,38^. We found that a measure of non-Gaussianity, the Anderson-Darling statistic^39^, was markedly reduced by SIQReg (median reduction of 85.8%, **Fig. 3d**). We also found SIQReg reduced heteroscedasticity induced by mean–variance coupling^40^ (median reduction of 85.3%, **Fig. S7**). These improvements are evident in the phenotype distributions; even testosterone, which is bimodal due to sex differences, more closely resembled a mixture of Gaussians on the SIQReg scale (**Extended Data Fig. 1**). Because SIQReg does not directly optimize for normality or homoscedasticity, these improvements provide independent evidence that SIQReg recovers a more appropriate scale.

### SIQReg removes substantial scale-induced genetic heterogeneity

Having established that the SIQReg scale is robust and reduces residual heterogeneity, we asked how it impacted downstream analyses of non-additive genetic effects, i.e., genetic heterogeneity. We evaluated genetic heterogeneity at three levels of genetic architecture: PGS tests using quantile regression and GxE; gene-level tests using imputed gene expression and quantile regression; and single-variant vQTL tests.

We found that 27% of PGS-trait pairs showed quantile-dependent heterogeneity on the default scale (239/900 had p < 0.05/36, adjusted for 36 PGS, **Fig. 4a**). Strikingly, over a third of these signals vanished on the SIQReg scale (34%). For example, the effect of the height PGS on height seemingly increases with height on the default scale, but its effect is additive on the SIQReg scale (**Fig. 4b)**. SIQReg learns a log-like scale for height (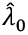 = −0.021, *SE* = 0.047), which is consistent with recent arguments that height is log-additive^26,32^ but inconsistent with standard practice^41^. In contrast, the BMI PGS remains heterogeneous after SIQReg transformation, with greater effects on individuals with higher BMI; this is consistent with twin-based heritability estimates increasing with BMI^42^ and with genetic correlations across BMI strata below one^43^. Similar conclusions held when we replaced our statistical test of heterogeneity across quantiles with a more qualitative measure of “non-equivalence”^19^ (**Fig. S8**). Overall, SIQReg can remove likely-spurious PGS heterogeneity while preserving likely-meaningful PGS heterogeneity.

**Figure 4.**
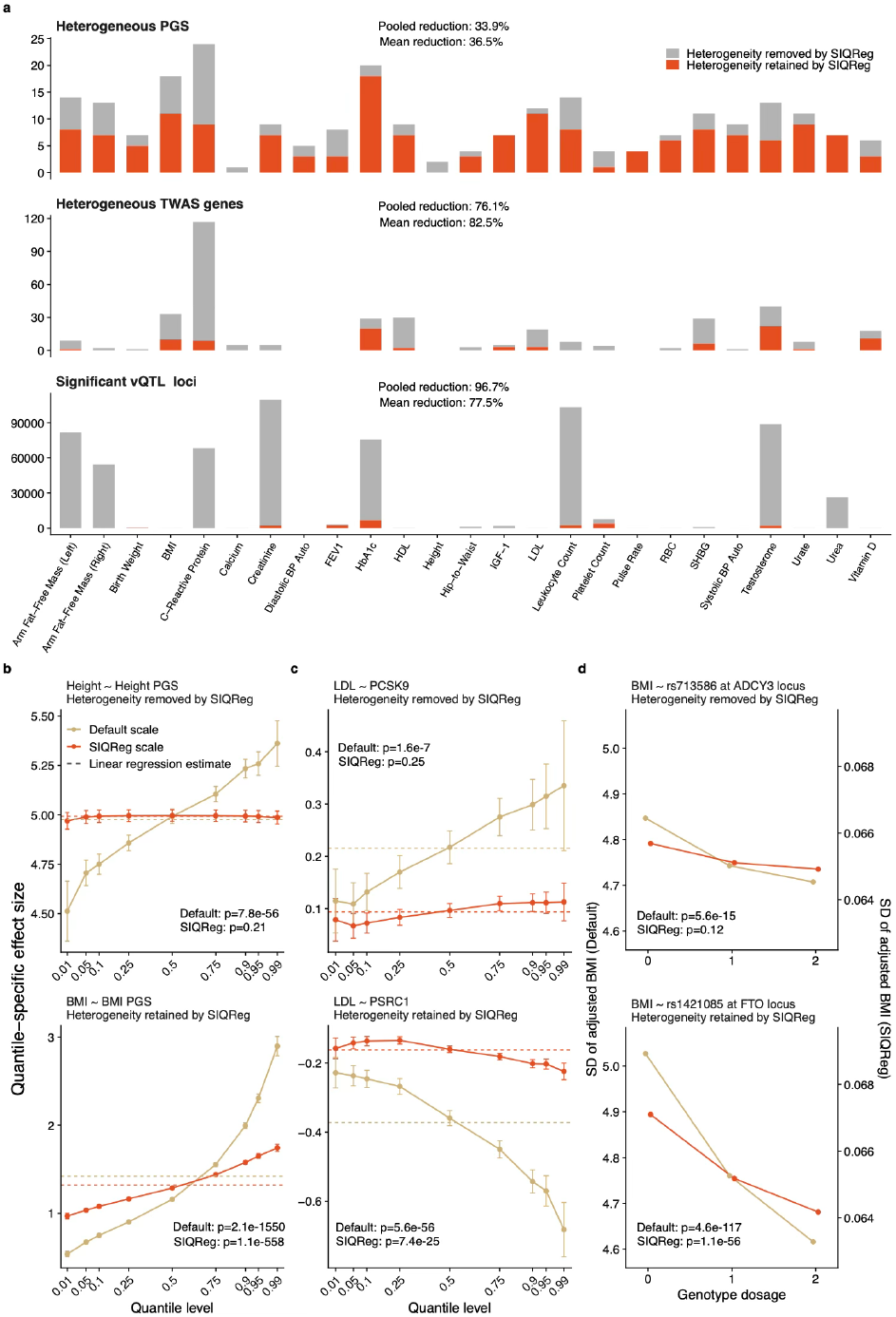
SIQReg improves inference on genetic heterogeneity. (a) Across 25 complex traits, the numbers of heterogeneous PGS, genes, and variants are shown on the default and SIQReg scales (only a few heterogeneous signals newly arise on the SIQReg scale and are shown in **Table S2-4**). We test variant-level heterogeneity with vQTL rather than quantile regression due to the computational cost of genome-wide quantile regression^20^. (b) The height PGS varies across quantiles on the default scale, but this vanishes on the SIQReg scale; conversely, the BMI PGS shows quantile-dependent effects on both scales. (c) *PCSK9* heterogeneity for LDL is removed by SIQReg, whereas *PSRC1* heterogeneity persists. (d) The vQTL effect of rs713586 on BMI is removed by SIQReg, whereas the vQTL effect of rs1421085 persists.

We next evaluated SIQReg’s impact on PGSxE tests, a widely used form of genetic interaction^14^. Using sex as an “E”, we found that 20.3% of significant PGSxSex signals on the default scale vanished after SIQReg transformation (30/148, p < 0.05/36, **Fig. S9**). The PGS-trait pairs that lost PGSxSex signals were enriched in PGS-trait pairs that lost quantile-dependent signals (*p* = 0.0058, **Fig. S10**), highlighting the connection between these heterogeneity tests (**Note S1.3**). We found similar results using age, smoking status, alcohol intake frequency, and statin use as “E” in PGSxE tests (5.8%, 16.7%, 5.3%, and 11.0%, respectively, **Fig. S11**).

Second, at the gene level, transcriptome-wide association study (TWAS) tests of quantile heterogeneity revealed similar patterns: on average across phenotypes, 76% of heterogeneous TWAS genes were removed by SIQReg transformation (**Fig. 4a**). We highlight two genes with distinct patterns of effects on LDL cholesterol (**Fig. 4c**). *PCSK9* has a heterogeneous default-scale effect that is removed by SIQReg, consistent with evidence that *PCSK9* inhibitors are broadly effective across patient populations^44^. In contrast, *PSRC1* has quantile-dependent effects on both default and SIQReg scales, suggesting its heterogeneity is scale-independent. We hypothesized that statin use explained this discrepancy^45^, so we repeated the analysis restricted to non-users. Indeed, we found that *PCSK9* was homogeneous on the default scale after removing statin users (*p* = 0.84), whereas *PSRC1* remained heterogeneous (*p* = 8.5e − 7). This suggests that SIQReg appropriately removes confounding by statins while preserving more biologically meaningful heterogeneity.

Third, we investigated single-variant-level heterogeneity using vQTL tests (**Fig. 4a**). We found that SIQReg eliminated the vast majority of vQTL across traits (97%). For 12/24 traits, over 95% of the vQTL vanished after SIQReg transformation. This is expected based on our finding that default scales induce substantial mean–variance coupling that is removed by SIQReg. Nonetheless, SIQReg preserved plausible vQTL, such as the well-known effect of the *FTO* locus for BMI^46^ that is implicated in diverse biological pathways including adipocyte thermogenesis^47,48^ and satiety^49^ (**Fig. 4d**). We found that default-scale vQTL were severely inflated genome-wide, whereas the SIQReg scale gave calibrated tests (median *λ*_*GC*_ reduced 17%, **Fig. S12**), supporting the interpretation that SIQReg vQTL are meaningful whereas default vQTL are likely spurious.

Overall, SIQReg transformation substantially reduces genetic heterogeneity signals in multiple analyses while preserving biologically plausible forms of genetic heterogeneity.

### SIQReg improves genetic discovery

We showed that SIQReg removes default-scale heterogeneity signals. In principle, this could be explained by SIQReg identifying a better scale that is more additive, or a worse scale that simply has lower power to detect heterogeneity. To resolve this, we asked how SIQReg impacted power to detect additive signals.

We found that SIQReg increased the number of GWAS loci by 11.2% on average across traits, though the number slightly decreased for some traits (**Fig. 5a**). One example of a new GWAS signal from SIQReg is rs148008104 at the *STEAP1B* locus for C-reactive protein, which was previously reported in large-scale GWAS^50,51^ and is involved in metal-ion reduction^52^. This signal replicated across ancestry groups on the default scale (p < 0.05), showing how the SIQReg scale adds power to detect genuine additive signals (**Fig. 5b**).

**Figure 5.**
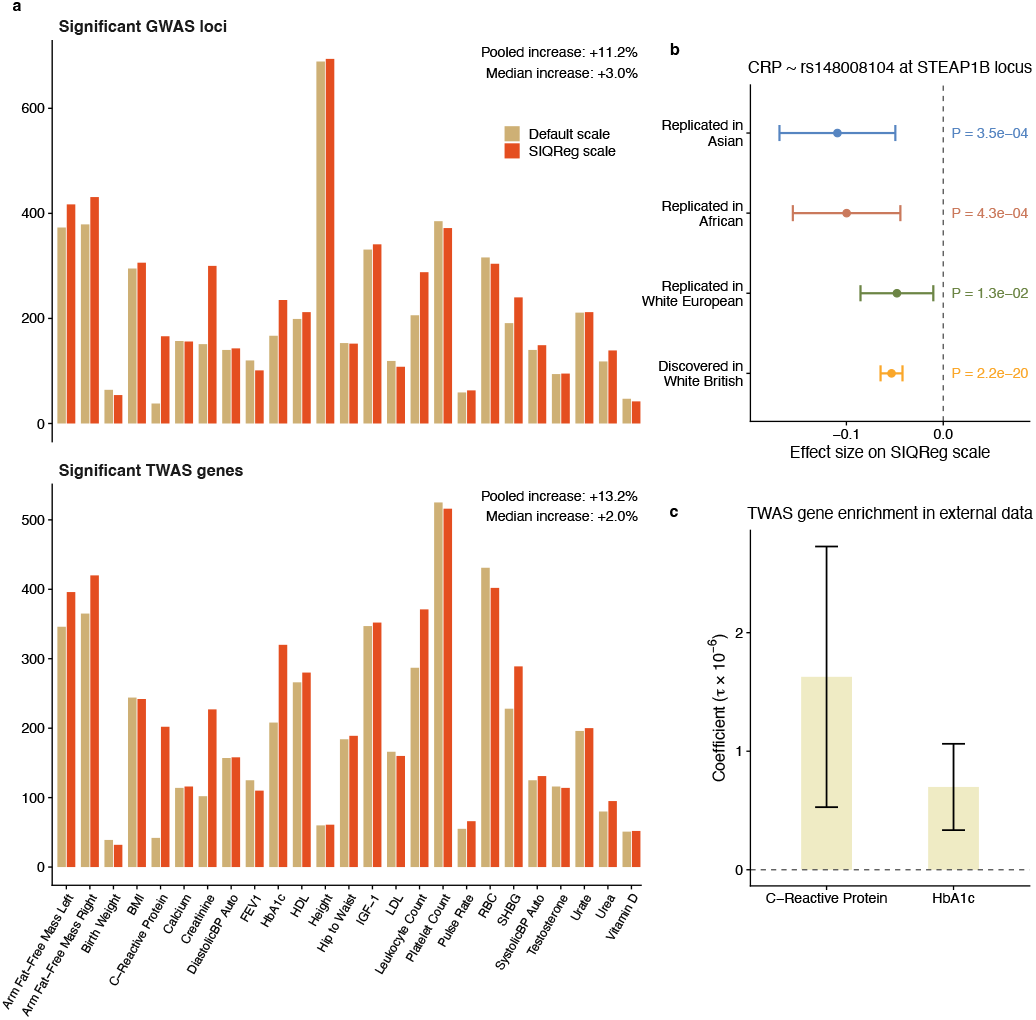
SIQReg improves genetic discovery power. (a) Number of significant GWAS loci (top) and TWAS genes (bottom) on the default and SIQReg scales across 25 complex traits in UK Biobank. (b) A novel GWAS variant identified by SIQReg transformation in White British, rs148008104, replicates in other UK Biobank ancestry groups on the default scale. (c) Two sets of novel TWAS genes identified on the SIQReg scale are enriched in GWAS heritability in external cohorts.

We found similar gains for TWAS, where SIQReg increased the number of TWAS hits by 13.2% across traits (**Fig. 5a**). Because these discoveries use European-ancestry TWAS weights, we did not validate in other UK Biobank ancestry groups. Instead, we used LDSC to test enrichment of the regions around newly associated genes in external GWAS for two traits (C-reactive protein and HbA1c), both of which replicated (*p* = 0.007, 0.002; **Fig. 5c**).

Finally, we found that SIQReg dramatically improved calibration in the “additive-variance” test, which jointly tests a variant’s additive and vQTL effects^40^. For example, for C-reactive protein, we found severe inflation on the default scale (*λ*_*GC*_ > 10) that was largely solved on the SIQReg scale (*λ*_*GC*_ = 1.32, **Extended Data Fig. 2**). This indicates that the nominal power gain in the “additive-variance” test is not generally reliable.

Overall, SIQReg improved additive discovery power, which is theoretically expected (**Note S1.4**) and suggests that the SIQReg scale is superior.

### SIQReg improves polygenic prediction

Having established that analyses on the SIQReg scale improve GWAS discovery, we asked if this yielded downstream improvements in PGS (**Methods**). We assessed PGS prediction accuracy using Spearman *R*^2^ rather than Pearson *R*^2^ because it is less scale-dependent. We found that SIQReg-PGS were generally superior: they significantly improved *R*^2^ for 13/25 traits (p < 0.05) and on average increased *R*^2^ by 10.3% (95% CI: 8.1%-12.5%, **Fig. 6a**). Nonetheless, the default scale significantly outperformed SIQReg for 3/25 traits (p < 0.05).

**Figure 6.**
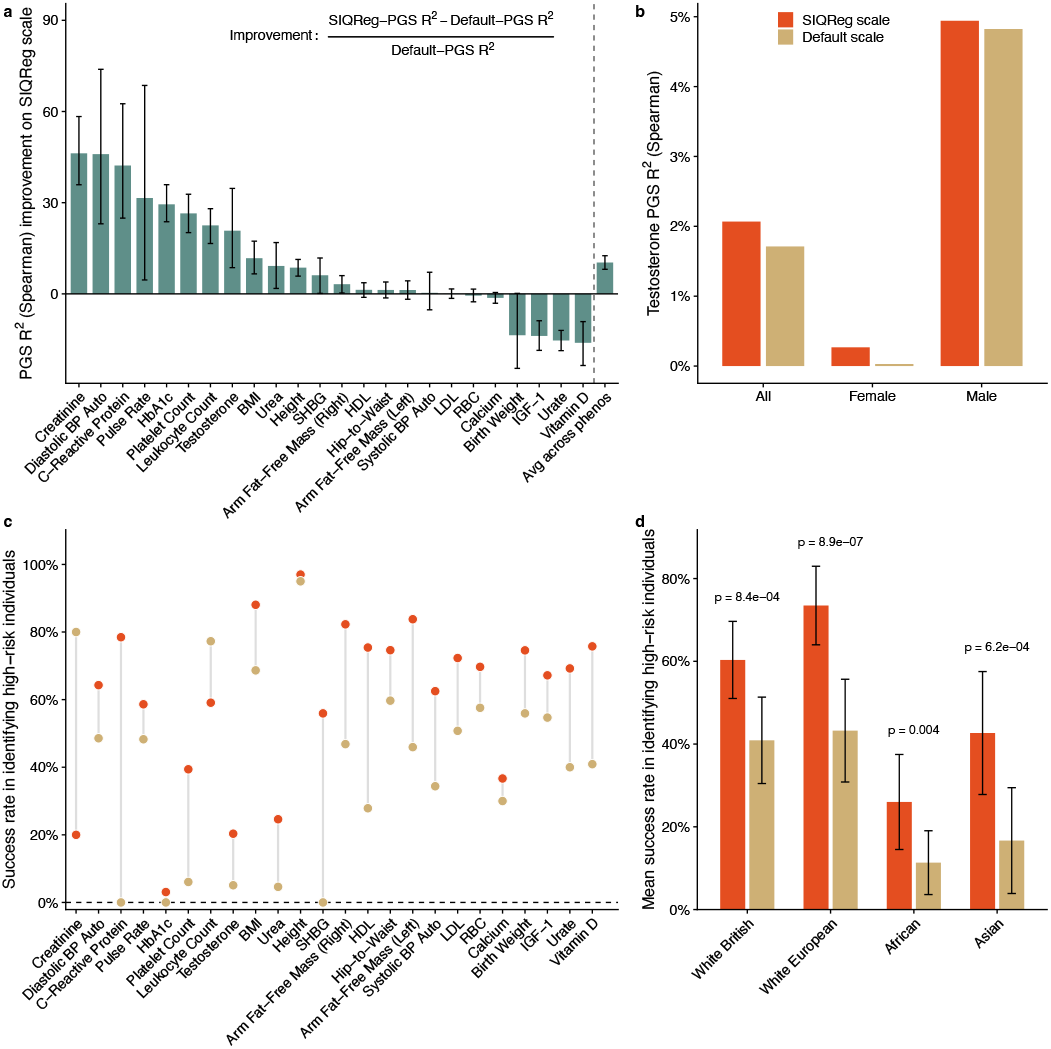
SIQReg improves PGS prediction accuracy and high-risk individual identification. (a) Percentage change in prediction accuracy of PGS constructed on the SIQReg scale relative to the default scale; positive values indicate improvement after SIQReg transformation. Prediction accuracy was measured using Spearman correlation-based *R*^2^. (b) Testosterone PGS prediction accuracy on the default and SIQReg scales calculated in all individuals and separately in females and males. (c) PredInterval success rate for identifying high-risk individuals (top 0.1%) using PGS on the default and SIQReg scales. (d) Mean success rate for high-risk identification across traits in four UK Biobank ancestry groups.

The case of testosterone illustrates a clear benefit of the data-driven SIQReg scale over simply comparing the default and log scales. Previously, we described a trade-off where log-scale PGS (*λ*_0_ = 0) performed better for females^26^ while default-scale PGS (*λ*_0_ = 1) performed better for males; here, we found that SIQReg strikes a balance (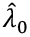 = 0.41) such that it simultaneously improves PGS accuracy for both sexes (**Fig. 6b**).

We next asked if SIQReg improved detection of high-risk individuals using PredInterval^7^ to predict which individuals lie in the top 0.1% of the phenotype distribution. We found that SIQReg-PGS produced calibrated prediction intervals (**Fig. S13**) and identified 1.5-fold more high-risk individuals across traits (**Fig. 6c, Fig. S14**). Consistent results held when using a top 1% threshold (**Fig. S15-16**). We validated these results by comparing results across three additional ancestry groups in UK Biobank; in each case, SIQReg achieved significantly higher mean success rates (**Fig. 6d**).

Interestingly, SIQReg even improved tail-risk prediction for the three traits where SIQReg-PGS reduced the Spearman *R*^2^ (Urate, IGF-1, Vitamin D). To understand why, we performed PredInterval on the default scale with the SIQReg-PGS and vice-versa. We found that the gains primarily resulted from performing PredInterval on the SIQReg scale, rather than building using the SIQReg-scale GWAS (**Extended Data Fig. 3, Fig. S17-18**), indicating that the gain comes not from better genetic effect estimation but from the scale on which prediction is performed. This is expected because the SIQReg scale produces more Gaussian and homoscedastic noise terms, which matter most for identifying high-risk individuals.

## Discussion

The choice of phenotype measurement scale has been discussed for at least a century^34–36^. While some studies carefully consider, stress-test, and report this choice; and other studies explicitly choose a scale based on interpretability or known underlying biology; most studies implicitly choose whatever default scale is provided. This has become a major problem due to the size of biobank-based GWAS, which have sufficient power to detect subtle forms of model misspecification^26,53^ and often agnostically analyze many traits in parallel^54,55^. Here, we have proposed a simple, automatic, and general solution to this problem called SIQReg, which learns a trait-specific scale by minimizing heterogeneity across quantiles of the phenotype distribution.

Our results inform the nature of non-additive genetic effects in complex traits. The field has been dominated by additive models, although it has been recognized since Fisher’s foundational work^56^ that scale transformation may be needed. Our work shows that standard tests of non-additivity are highly prone to detect signals that can be eliminated by scale transformation, including GxE tests, vQTL tests, and quantile-heterogeneity tests. At the same time, we found that SIQReg preserves many non-additive signals, including biologically plausible signals such as the vQTL effects of *FTO* on BMI. We also found that the SIQReg scale improves power to detect additive signals and replicates across ancestry groups, further supporting its validity. Together, our results suggest a middle ground on non-additive genetic effects in complex traits: many are scale-dependent artifacts, yet many others are genuine scale-independent signals.

Our consistent finding that *λ*_0_ < 1 across 25 traits addresses a classic question in quantitative genetics: are phenotypes additive or multiplicative^2,57^? Our estimates of *λ*_0_ indicate a mix of additive and multiplicative effects, such that neither default nor log scales are perfectly additive. We developed a mathematical model that can explain this finding, in which each variant acts through a mix of additive and multiplicative pathways (**Note S1.5**). Unlike our null model in SIQReg, where all effects are additive on some latent scale, this more flexible model allows different biological pathways to act on different scales.

SIQReg distinguishes traits that become additive after transformation from those that remain non-additive across scales, such as height and BMI, respectively. Height has long been understood to be approximately log-additive^32^; SIQReg directly confirms this because it estimates a near-log transformation that, in turn, eliminates all non-additive signals detected on the default scale. By contrast, BMI is a ratio of heritable phenotypes, suggesting that no monotone transformation will recover additivity^58^. SIQReg also supports this because it only eliminates some of the non-additive signals detected on the default scale. In particular, we find that individuals at the upper tail of the BMI distribution have higher PGS than expected, consistent with their qualitative differences in comorbidity burden^59,60^ and differential response to certain treatments, such as setmelanotide being effective only for individuals with rare melanocortin pathway variants^61,62^.

This work has several limitations. First, SIQReg learns a monotone transformation within the Box-Cox family. Extending to more flexible transformations may be required for phenotypes with complicated distributions, such as bounded phenotypes. Second, likelihood-based alternatives such as WarpedLMM are more powerful than our semiparametric approach when their assumptions hold, but we found that they are highly sensitive to non-Gaussian noise. Third, we have assumed continuous phenotypes, but similar considerations apply to the choice of link function for ordinal phenotypes, such as never/current/ever smoker, and binary case/control phenotypes^63,64^; the primary challenge of extending SIQReg to these phenotypes is power. Fourth, SIQReg estimates the optimal scale using covariates that are themselves measured on default scales; future work should consider joint transformations of covariates and phenotypes. Fifth, we have relied on nongenetic covariates to estimate the scale for power. Although our results were generally robust to using PGS instead of covariates, in principle, variance component models could be a powerful approach to directly optimize phenotype scale for genetic discovery. Finally, while SIQReg distinguishes scale-dependent from scale-independent heterogeneity, the former may still be biologically meaningful and warrant further investigation. Nonetheless, we expect that characterizing scale-dependence will generally add insights into studies of non-additive genetic effects.

Overall, SIQReg is a practical and robust approach to optimize phenotype scale that improves genetic analyses of complex traits.

## Methods

### Quantile regression

Instead of estimating effects on the conditional mean of the phenotype, quantile regression focuses on its conditional quantiles, characterizing the entire conditional distribution of a phenotype of interest. Let *y*_*i*._ denote the default-scale phenotype, *G*_*i*._ denote a genetic factor, *C*_*i*._ denote a vector of covariates and 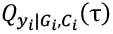 denote the *τ*-th conditional quantile function of *y*_*i*._. Then the linear quantile regression model for a given quantile level *τ* ∈ (0,1) is

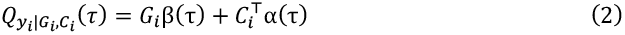

where *β*(*τ*) and *α*(*τ*) are quantile-specific coefficients. The quantile regression estimator minimizes the check function:

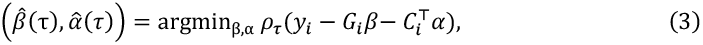

where *ρ*_*τ*_ (*u*) *= u*(*τ* − *I*(*u* < 0)) is the asymmetric absolute loss function that weights positive and negative residuals by *τ* and 1 − *τ*, respectively.

Classical quantile regression involves minimizing a piecewise linear check loss function, which can be reformulated as a linear program. Standard interior-point algorithms have computational complexity *O*(*pn*^*1+c*^ log *n*) for some *c* ∈ (0, 1/2) with *p* being the number of coefficients and *n* being the number of individuals, which becomes prohibitive for biobank-scale analyses with hundreds of thousands of individuals and dozens of predictors across multiple quantile levels. To achieve scalable computation, we implement all quantile regressions using the conquer (convolution-type smoothed quantile regression) framework^65^, which replaces the non-differentiable check function *ρ*_*τ*_ (*u*) with a smoothed surrogate obtained by convolution with a kernel function:

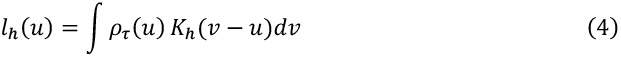

where *K*_*h*_(*v − u*) *= h*^*-1*^*K*(*u*/*h*) is a scaled kernel function with a positive bandwidth *h*. When *K*(·) is a non-negative kernel (we use the Gaussian kernel throughout), the smoothed loss is globally convex and twice continuously differentiable, enabling efficient gradient-based optimization with complexity *O*(*pn*) per iteration. Under mild regularity conditions, the resulting smoothed estimator achieves the same 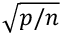 convergence rate as classical quantile regression when the bandwidth satisfies *h* ≍ (*p* + log*n*)/*n*^γ^ for γ ∈ [1/4,1/2]. We use the recommended bandwidth γ = 2/5.

For testing the null hypothesis of genetic heterogeneity *H*_0_: *β*(*τ*_1_) = … = *β*(*τ*_k_), which tests whether the genetic effect is quantile-dependent, let 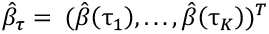 denote the vector of estimated genetic effects, and let Σ_*β*_ denote its covariance matrix, the test statistic is:

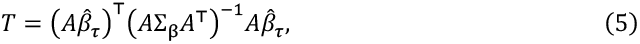

where *A* is a (*K*− 1) × *K* matrix with row (1, −1,0, …, 0), (0,1, −1, …, 0), …, (0, …, 0,1, −1). Under the null, this test statistic asymptotically follows a chi-squared distribution with degrees of freedom *K*− 1.

### SIQReg

Scale-independent quantile regression (SIQReg) learns a trait-specific monotone transformation that recovers a latent scale with minimal scale-induced heterogeneity. We assume that there exists a latent phenotype *y*\* such that:

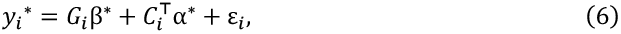

where *β*\* and *α*\* are true effects on the latent scale and ε is the error term. Under the assumption that covariates are independent of the error, the conditional quantile function satisfies:

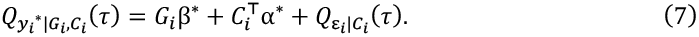

The default-scale phenotype *y* is linked to *y*\* through a monotone transformation 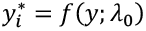 where *f*(·) belongs to a parametric family indexed by *λ*_0_. In this paper, we focus on the Box-Cox transformation family:

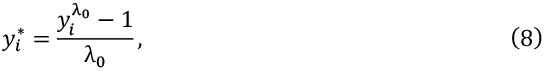

which reduces to the log transformation in the limit *λ*_0_ → 0. The key insight underlying SIQReg is that on the correct latent scale, quantile regression coefficients should be constant across quantile levels. Conversely, heterogeneous coefficients for covariates across quantiles are expected on the incorrect scale. Therefore, SIQReg seeks the most-additive scale by estimating *λ*_0_ to minimize coefficient variation across quantiles.

Specifically, for a candidate scale *λ*, SIQReg first fits quantile regression into 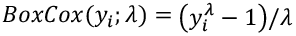 quantile levels *τ*_1_ < … < *τ*_*k’*_ yielding quantile-specific coefficients for *j*-th covariate: 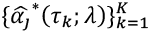. To assess heterogeneity across quantiles, let 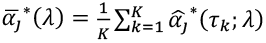 and 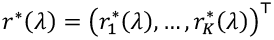 where 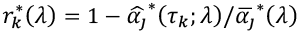 and Σ(*λ*) is its covariance matrix. Since *λ*_0_ is an intrinsic property of a trait and does not depend on the choice of covariate, we obtain estimates for each covariate and then meta-analyze. SIQReg estimates *λ*_0_ by minimizing a Wald statistic measuring quantile-dependent heterogeneity:

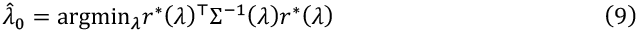

To construct the confidence interval, we adopt a subsampling strategy where we split the full sample into *R* disjoint subsamples and use the standard deviation of subsample estimates^66^. After estimating 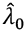, SIQReg fits quantile regression on the transformed scale

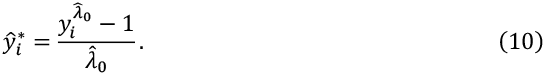

In practice, we use *K* = 9 quantile levels *τ* ∈ {0.1, 0.2, …, 0.9} for inferring *λ*_0_ to reduce noise and *τ* ∈ {0.01, 0.05, 0.1, 0.25, 0.5, 0.75, 0.9, 0.95, 0.99} for testing genetic heterogeneity, where we use the bootstrap to estimate the covariance matrix Σ(*λ*) and Σ_β_. We estimate *λ*_0_ using non-genetic covariates as input by searching over candidates in [−5, 5] as they typically explain more phenotypic variance than any single genetic predictor. We implement SIQReg one-covariate-at-a-time and then meta-analyze the 43 results to obtain 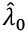. Because the transformation parameter is a property of the phenotype measurement scale rather than specific predictors, *λ*_0_ estimated from covariates generalizes to genetic analyses.

### WarpedLMM

We compared SIQReg to WarpedLMM, a likelihood-based approach that estimates phenotype transformation. We adopt the term WarpedLMM because our implementation shares its core principle of maximizing a Gaussian likelihood, though the original WarpedLMM uses the linear mixed model likelihood and considers other classes of transformations. Specifically, WarpedLMM infers the transformation *λ*_0_ by maximizing the log-likelihood plus a Jacobian term:

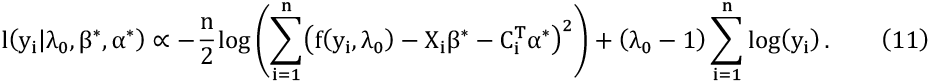

The key difference is that WarpedLMM assumes Gaussian errors on the latent scale, which can lead to biased estimates when this assumption is violated (e.g., heavy-tailed errors). SIQReg, as a semiparametric approach, makes no distributional assumptions, instead relying only on the quantile regression framework.

### Simulations

We conducted simulations to evaluate SIQReg’s ability to (1) recover the true latent scale, (2) distinguish scale-induced artifacts from genuine heterogeneity on the latent scale, and (3) enhance power for association analyses. Simulations generated default-scale phenotypes from a latent additive model *y*\* = *β*_0_ + *β*_*G*_ · *G* + *α* · *C* + ε, where the default-scale phenotype was obtained via the inverse Box-Cox transformation 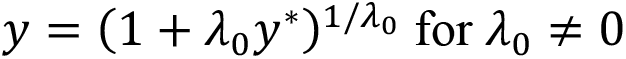 and *y* = exp(*y*\*) for *λ*_0_ = 0. All simulation settings were repeated across 1,000 replicates.

To evaluate recovery of the true latent scale, we simulated phenotypes under a range of *λ*_0_ values and sample sizes. We considered heavy-tailed errors drawn from a Student’s t-distribution to assess robustness to non-Gaussian errors, and in a separate experiment varied the degree of tail heaviness while holding the latent scale fixed. For comparison, we also estimated *λ*_0_ using the Gaussian likelihood framework underlying WarpedLMM.

To evaluate calibration of the heterogeneity test, we simulated phenotypes under the null of no latent genetic heterogeneity and compared heterogeneity detection on the default, log, RINT, and SIQReg scales across a range of true *λ*_0_ values. To evaluate power, we fixed the latent scale and introduced heteroscedastic errors of the form *ε*_*i*_ = exp(*γ* · *G*_*i*_) · *z*_*i*_ with *z*_*i*_ ~ *N*(0,1), so that quantile-dependent genetic heterogeneity truly existed on the latent scale and increased with γ.

To evaluate whether SIQReg improves association testing, we fixed the latent scale and simulated phenotypes with Student’s t errors while varying the additive genetic effect size. We then compared the power of linear regression on the default, log, and SIQReg scales. RINT was not included in power comparisons because it was severely miscalibrated in our simulations.

### UK Biobank analyses

We analyzed 25 quantitative traits (**Table S5**) in UK Biobank^67^. Our primary analyses used unrelated individuals of White British ancestry (n = 335,461). For cross-ancestry validation, we additionally analyzed individuals of White European (non-British; n = 24,951), African (n = 7,488), and Asian (n = 2,459) ancestry. SIQReg estimation used 43 covariates: age, age^2^, sex, and the first 40 genetic principal components.

### GWAS and PGS construction

We implemented GWAS and PGS construction following the pipeline in Costantino et al.^26^. Briefly, GWAS were run using PLINK2^68^ on 6,539,458 SNPs passing standard quality filters in a 90% training set of White British individuals, with age, sex, and 10 genetic principal components as covariates. Significant loci were identified at p < 5 × 10^−8^ after LD clumping (*R*^2^ < 0.1, 250 kb window). PGS were constructed by clump-and-threshold with p-value thresholds ranging from 5 × 10^−8^ to 0.5, selecting the optimal threshold by *R*^2^ on the GWAS scale.

PGS were evaluated in 24,950 held-out White European individuals. For default-scale GWAS, we used the following model:

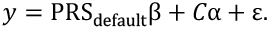

For SIQReg-scale GWAS, we used an alternative model to adjust for this change in scale.

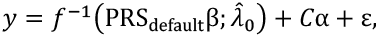

and analogously for the log scale.

### PGS heterogeneity and PGSxE

PGS quantile-dependent heterogeneity was tested using externally trained polygenic scores^69^ (**Table S6**). For PGS×E interaction analysis, we fitted the model from Costantino et al.^26^:

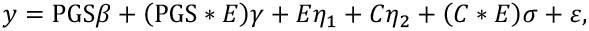

across five environmental variables (sex, age, statin use, smoking status, alcohol intake frequency), with environmental definitions as in Costantino et al.^26^ (**Table S7**). Statin usage labels followed Sadowski et al.^45^. The covariate–environment interaction term controls for confounding^70^.

### TWAS analysis

We performed transcriptome-wide association studies using PrediXcan with gene expression prediction models trained in GTEx v8^71^. We used the MASHR prediction models^4,72,73^ to impute gene expression. For each trait, we chose to use a tissue with plausible biological relevance (e.g., whole blood for LDL; **Table S8**) and additionally performed analyses across all tissues with Bonferroni correction for the number of genes and tissues tested.

### vQTL analysis and mean-variance coupling assessment

We identified vQTL, estimated mean-variance coupling effects, and implemented the Additive-Variance (AV) effect tests using the heteroscedastic linear mixed model (HLMM)^40^. We applied HLMM to both default and SIQReg-transformed phenotypes. Genome-wide significance was defined at a p-value threshold of 5 × 10^−8^.

### External data validation

For C-reactive protein, we obtained GWAS summary statistics from the CHARGE Inflammation Working Group meta-analysis^74^, which includes approximately 200,000 individuals of European ancestry from multiple cohorts excluding UK Biobank. We used these summary statistics to test enrichment of newly significant TWAS genes.

For HbA1c, we obtained GWAS summary statistics from the MAGIC consortium^75^, which includes approximately 150,000 individuals of European ancestry from multiple cohorts excluding UK Biobank. We used these summary statistics for similar validation analyses.

For gene enrichment analysis, we tested whether TWAS genes newly identified as significant on the SIQReg scale showed enrichment for association signals in external datasets. For each newly significant gene, we used LD score regression to test enrichment with baselineLD model v2.3^76^.

### PGS Prediction Interval

We constructed calibrated prediction intervals for PGS-based phenotype prediction using PredInterval^7^. Briefly, PredInterval uses K-fold cross-validation to compute PGS-based prediction residuals on a calibration set, then constructs prediction intervals for held-out individuals with approximately valid coverage at a specified confidence level, without distributional assumptions.

We split White British individuals into a training pool (80%) and a held-out test set (20%). Within the training pool, we performed 5-fold cross-validation: for each fold, GWAS was run on the remaining four folds with the same quality control, covariate specification and PGS construction as above. Cross-validated PGS were computed for both the held-out fold (for residual calibration) and the test set (for final evaluation). For cross-ancestry evaluation, we applied White British PGS weights to Asian, African, and White European non-British individuals in UK Biobank, using the *λ*_0_ estimated in White British to transform phenotypes in each group. Each ancestry group was split into a calibration set and a test set (50/50 for Asian due to small sample size; 70/30 for African and White European). Prediction intervals were then generated at the 95% confidence level.

This procedure was run independently on default and SIQReg phenotype scales. We report the success rate of identifying high-risk individuals, defined as the proportion of individuals with true phenotypic values in the top 1% and 0.1% that are identified by PredInterval. Statistical significance was assessed using two-sided paired *t*-tests across traits.

To determine whether improvements from SIQReg transformation are driven by improved PGS accuracy or by better residual calibration on the transformed scale, we performed a 2 × 2 factorial decomposition. The design crossed PGS source (default-scale or SIQReg-scale GWAS) with residual calibration scale (default or SIQReg PredInterval), with back-transformation applied to align PGS and phenotype scales as above.

## Data availability

The individual-level genotype and phenotype data are available to approved researchers through the UK Biobank web portal at https://www.ukbiobank.ac.uk/. The prediction weight used to impute gene expression can be downloaded at https://predictdb.org/post/2021/07/21/gtex-v8-models-on-eqtl-and-sqtl/. The summary statistics of the CHARGE C-reactive protein and MAGIC HbA1c GWAS used in this study are publicly available from the IEU open GWAS project accession code ieu-b-35 (https://opengwas.io/datasets/ieu-b-35), and MAGIC official site (https://magicinvestigators.org/downloads/index.html). BaselineLD v2.3 annotations can be found at https://zenodo.org/records/10515792. Plink v2.0 can be found at https://www.cog-genomics.org/plink/2.0/. S-LDSC software can be found at https://github.com/bulik/ldsc. Conquer software can be found at https://github.com/WenxinZhou/conquer. HLMM software can be found at https://github.com/AlexTISYoung/hlmm. PredInterval software can be found at https://github.com/xuchang0201/PredInterval.

## Code availability

The SIQReg package and all code used for analyses are available on GitHub at https://github.com/zhenhong7/SIQReg.

## Extended Data

**Extended Data Fig. 1.**
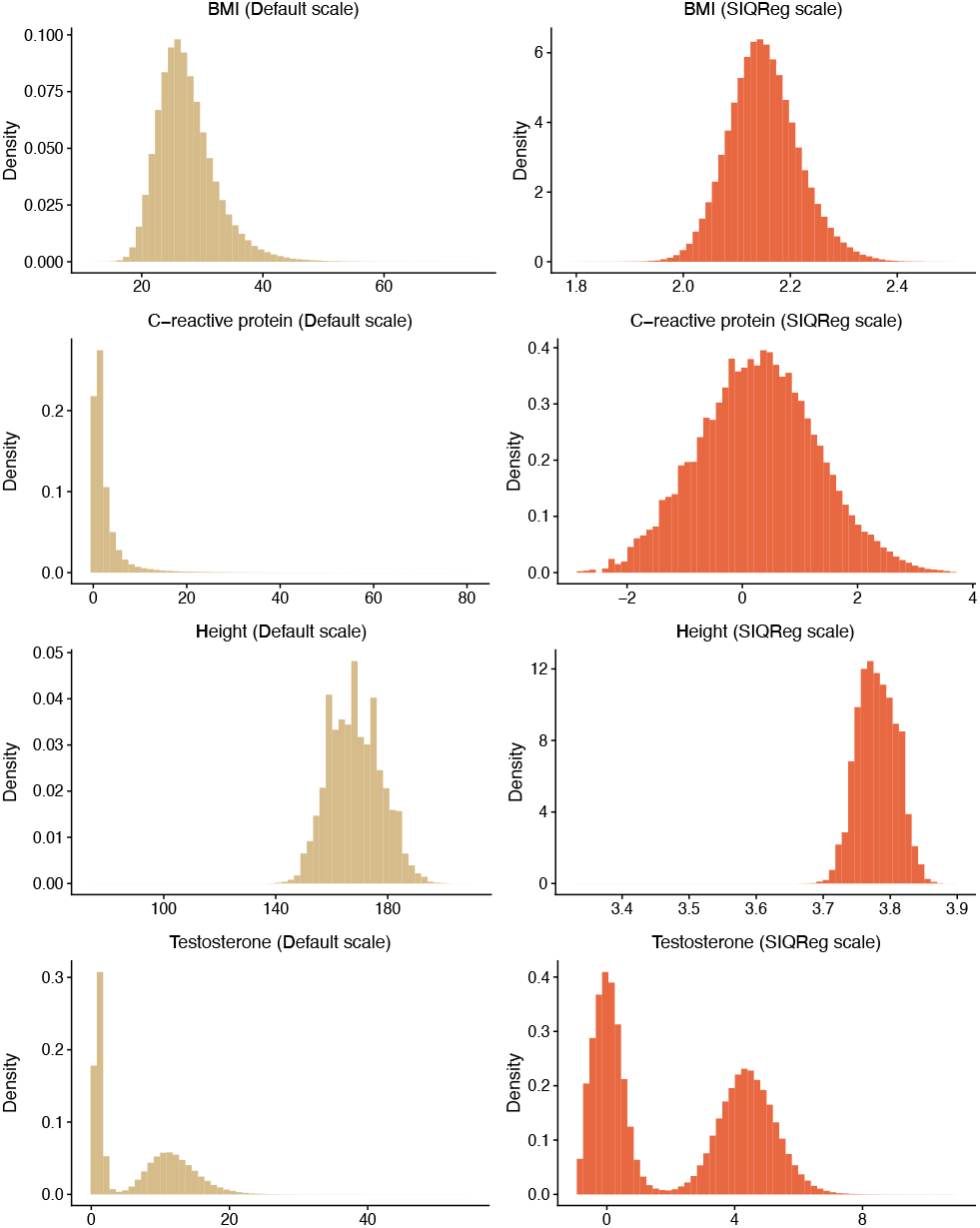
Distributions of BMI, C-Reactive protein, height and testosterone on the default measurement scale and SIQReg scale.

**Extended Data Fig. 2.**
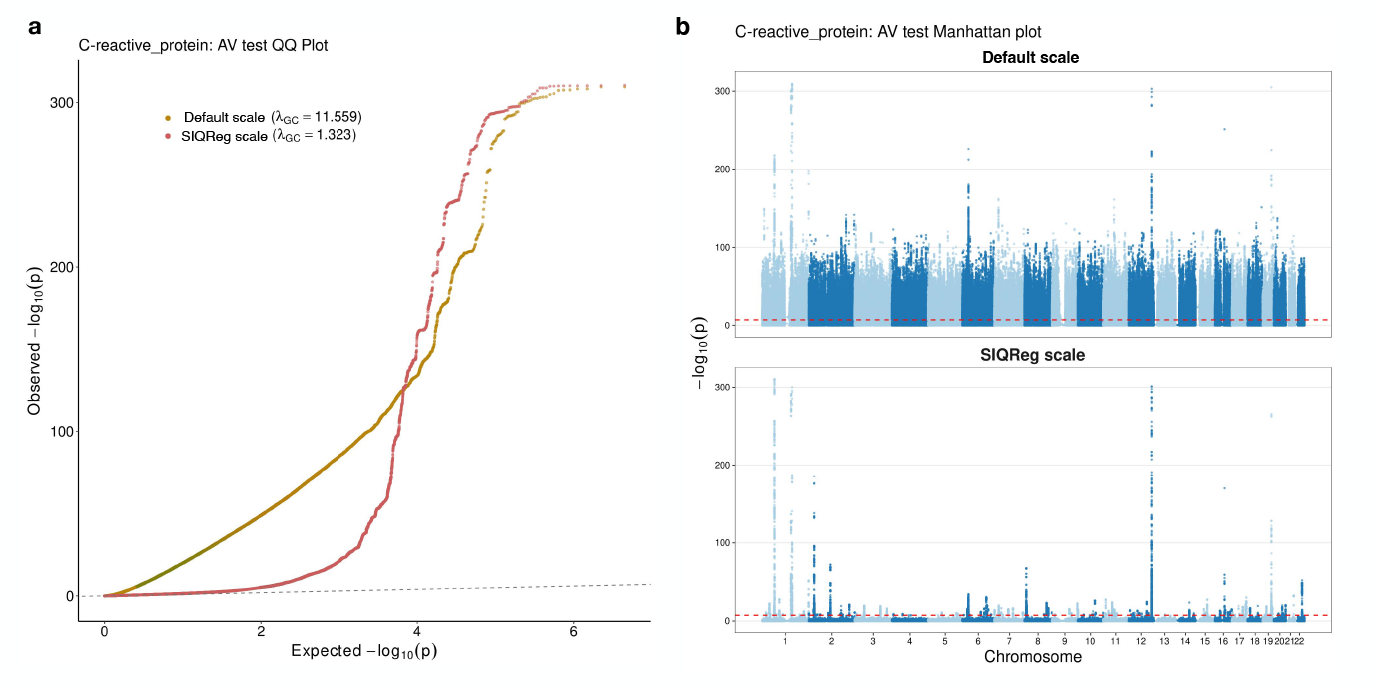
(a) QQ plot of additive variance test statistics for C-reactive protein, showing severe inflation on the default scale (*λ*_*GC*_ = 11.559) that is corrected on the SIQReg scale (*λ*_*GC*_ = 1.323). (b) Manhattan plots of additive variance tests for C-reactive protein on the default scale (top) and SIQReg scale (bottom).

**Extended Data Fig. 3.**
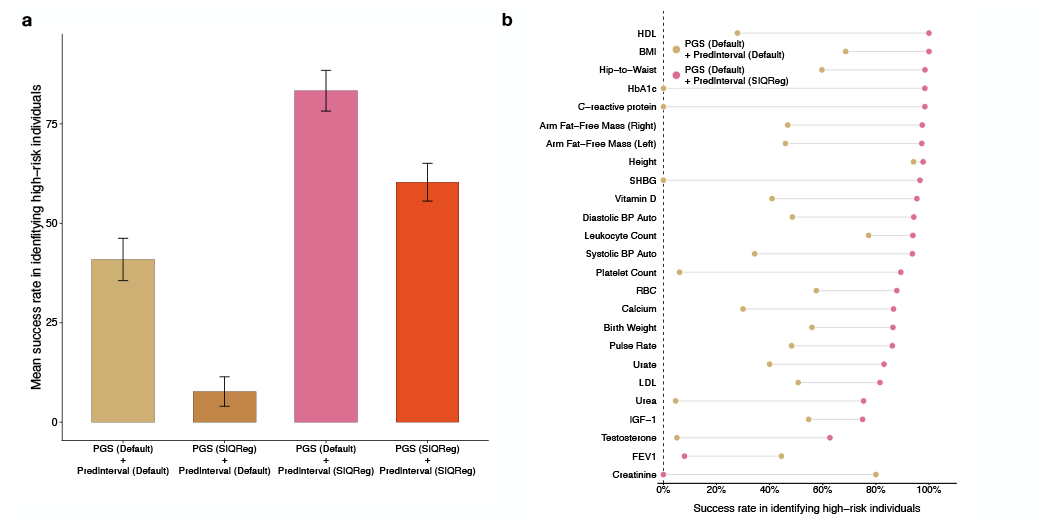
(a) Mean success rate across 25 traits for identifying high-risk individuals (top 0.1%) under four combinations of phenotype scale used for PGS construction and prediction interval estimation: Default PGS + Default prediction interval, SIQReg PGS + Default prediction interval, Default PGS + SIQReg prediction interval, and SIQReg PGS + SIQReg prediction interval. (b) Trait-by-trait comparison between Default PGS + Default prediction interval and Default PGS + SIQReg prediction interval. Holding the PGS fixed on the default scale shows that part of the improvement in tail identification is attributable specifically to performing prediction on the SIQReg scale.

## Acknowledgements

This research has been conducted using the UK Biobank Resource under Application Number 89052. A.D. is supported by R35GM150822. M.C. is supported by Fonds de Recherche du Québec Santé. We thank the participants in UK Biobank for making this study possible. Compute resources for this study were provided by the Randi HPC Cluster maintained by the Center for Research Informatics (CRI) at University of Chicago. The Center for Research Informatics is funded by the Biological Sciences Division and the Institute for Translational Medicine/CTSA (NIH UL1TR002389) at the University of Chicago. We thank Sriram Sankararaman (UCLA), Jonathan Flint (UCLA) and Iain Mathieson (University of Pennsylvania) for helpful feedback.

## Author contributions

Z.H. and A.D. developed the statistical methodology, conducted the analyses, and wrote the manuscript. M.C. provided critical suggestions and feedback on the manuscript.

