## Supplementary Information for "Optimizing phenotype scale improves genetic analyses in large-scale biobanks"

**2 Supplementary Figures** **12**

### 1 Supplementary Note

#### 1.1 Default phenotype scale can produce spurious nonadditive genetic effects

Researchers commonly analyze phenotypes on their default measurement scale $y_i$, for example values downloaded directly from UK Biobank. However, the default scale need not reflect how biological processes generate phenotypic variation. If the mapping from underlying biology to measured values is nonlinear, then analyzing $y_i$ directly can distort the architecture of genetic effects.

To formalize this, suppose that genetic effects are additive on some latent scale $y_i^*$:

$$y_i^* = G_i\beta^* + C_i^\top\alpha^* + \varepsilon_i, \tag{1.1}$$

where $G_i$ is the genetic factor of interest and $C_i$ is the vector of covariates. Furthermore, the latent scale is linked to the default scale by a monotone and nonlinear transformation $f(\cdot)$ parametrized by $\lambda_0$:

$$y^* = f(y; \lambda_0). \tag{1.2}$$

On the default scale, the marginal genetic effect is

$$\frac{\partial y}{\partial G} = \frac{\partial y}{\partial y^*} \cdot \frac{\partial y^*}{\partial G} = \beta^* \frac{df^{-1}}{dy^*}\Big|_{y^*=G\beta^*+C^\top\alpha^*+\varepsilon}. \tag{1.3}$$

Because $f(\cdot)$ is nonlinear, the term $\frac{df^{-1}}{dy^*}$ is not constant: it varies with the individual's phenotype level. This means the apparent genetic effect size on the default scale depends on where in the phenotype distribution the individual sits, producing spurious nonadditivity. For example, genetic effects that appear larger in the tails than near the mean, or vice versa.

Although the true biological scale is fundamentally unknown, it is possible to remove the scale-induced nonadditivity by estimating a transformation $f(y; \widehat{\lambda}_0)$. This does not claim to recover the true biological scale, but rather to find a working scale on which additive genetic models provide an adequate description of the data, thereby separating genuine genetic heterogeneity from artifacts of the measurement scale.

#### 1.2 SIQReg produces valid inference for the latent scale

##### 1.2.1 SIQReg estimation procedure

SIQReg aims at estimating $f(y; \widehat{\lambda}_0)$ with specifying $f(\cdot)$ to be Box-Cox transformation:

$$y_i^* = \frac{y_i^{\lambda_0} - 1}{\lambda_0}. \tag{1.4}$$

Under the latent scale, we have the conditional quantile of latent phenotype:

$$Q_{y_i^*|G_i,C_i}(\tau) = G_i\beta^* + C_i^\top\alpha^* + Q_{\varepsilon_i|G_i}(\tau). \tag{1.5}$$

under the assumption that $\varepsilon \perp\!\!\!\perp C$. Crucially, the coefficients $\alpha^*$ do not depend on $\tau$: they are constant across quantile levels. On any other scale with $\lambda \neq \lambda_0$, the nonlinear distortion causes these coefficients to vary with $\tau$, producing quantile-dependent effects that serve as a diagnostic for mis-specification of $\lambda$. SIQReg exploits this property by searching for the $\lambda_0$ that minimizes the variation of covariate effects across quantile levels. The procedure is summarized in Algorithm 1.

**Algorithm 1** SIQReg latent transformation estimation

**Require:** Default phenotype $\{y_i\}_{i=1}^n$, genetic factor $\{G_i\}_{i=1}^n$, covariates $\{C_i\}_{i=1}^n$, quantile levels $\{\tau_k\}_{k=1}^K$, candidate set $\Lambda$ for transformation parameter

**Ensure:** SIQReg transformation parameter $\widehat{\lambda}_0$

**Step 1: Quantile regression on candidate scales**

**for** each $\lambda \in \Lambda$ **do**

  Compute transformed phenotype: $y_i^*(\lambda) = (y_i^\lambda - 1)/\lambda$

  **for** each quantile level $\tau_k$, $k = 1, \ldots, K$ **do**

    Fit quantile regression of $y_i^*(\lambda)$ on $G_i$ and $C_i$:

    $Q_{y_i^*(\lambda)|G_i,C_i}(\tau_k) = G_i\,\widehat{\beta}^*(\tau_k;\lambda) + C_i^\top \widehat{\alpha}^*(\tau_k;\lambda)$

  **end for**

**end for**

**Step 2: Construct the sample relative deviation for $j$-th covariate**

**for** each $\lambda \in \Lambda$ **do**

  Compute mean: $\widehat{\bar{\alpha}}_j^*(\lambda) = \frac{1}{K}\sum_{k=1}^K \widehat{\alpha}_j^*(\tau_k;\lambda)$

  Compute relative deviation vector:

$$\widehat{r}^*(\lambda) = \left(\widehat{r}_1^*(\lambda), \ldots, \widehat{r}_K^*(\lambda)\right)^\top, \quad \text{where } \widehat{r}_k^*(\lambda) = 1 - \frac{\widehat{\alpha}_j^*(\tau_k;\lambda)}{\widehat{\bar{\alpha}_j^*(\lambda)}}$$

  Estimate $\widehat{\Sigma}_r(\lambda)$ with bootstrap, the $K \times K$ covariance matrix of $r^*(\lambda)$

**end for**

**Step 3: Estimation**

Solve:

$$\widehat{\lambda}_0 = \underset{\lambda\in\Lambda}{\operatorname{argmin}}\ \widehat{r}^*(\lambda)^\top\, \widehat{\Sigma}_r^{-1}(\lambda)\, \widehat{r}^*(\lambda)$$

**return** $\widehat{\lambda}_0$

##### 1.2.2 Practical normalization

When $\lambda_0 < 0$, the transformed values $y_i^*(\lambda)$ can be close to zero for all individuals, leading to numerically unstable quantile regression estimates. To avoid this, we rescale:

$$\widetilde{y}_i^*(\lambda) = \frac{y_i^*(\lambda) - \text{median}(y^*(\lambda))}{\text{MAD}(y^*(\lambda))}. \tag{1.6}$$

This does not change the minimizer because the relative deviation $r^*(\lambda)$ is invariant to affine transformations of the response. Specifically, for any $a \neq 0$ and $b$, quantile regression satisfies $\widehat{\alpha}(\tau; ay + b) = a\,\widehat{\alpha}(\tau; y)$, so that $r_k^*(\lambda) = 1 - \widehat{\alpha}_j^*(\tau_k; \lambda)/\bar{\alpha}_j^*(\lambda)$ is unchanged.

##### 1.2.3 Consistency of $\widehat{\lambda}_0$

We now show that SIQReg produces a consistent estimate of $\lambda_0$. Define the population criterion, a Mahalanobis distance:

$$MD_{\text{pop}}(\lambda) = r^*(\lambda)^\top \Sigma_r^{-1}(\lambda)\, r^*(\lambda), \tag{1.7}$$

where $r^*(\lambda) = (r_1^*(\lambda), \ldots, r_K^*(\lambda))^\top$ with $r_k^*(\lambda) = 1 - \alpha_j^*(\tau_k; \lambda)/\bar{\alpha}_j^*(\lambda)$. By construction, $MD_{\text{pop}}(\lambda_0) = 0$ since $\alpha_j^*(\tau_k; \lambda_0) = \alpha_j^*$ for all $k$. The sample analogue is

$$MD_n(\lambda) = \widehat{r}^*(\lambda)^\top \widehat{\Sigma}_r^{-1}(\lambda)\, \widehat{r}^*(\lambda). \tag{1.8}$$

Since $\widehat{\alpha}_j^*(\tau_k; \lambda) = \alpha_j^*(\tau_k; \lambda) + O_p(n^{-1/2})$ uniformly over $\Lambda$ (He et al., 2023), and assuming that $\inf_{\lambda \in \Lambda} |\bar{\alpha}_j^*(\lambda)| > 0$, under standard regularization conditions, we have

$$\sup_{\lambda \in \Lambda} |MD_n(\lambda) - MD_{\text{pop}}(\lambda)| \xrightarrow{p} 0, \tag{1.9}$$

and by the argmin continuous mapping theorem (Van der Vaart, 2000),

$$\widehat{\lambda}_0 = \text{argmin}_{\lambda \in \Lambda}\, MD_n(\lambda) \xrightarrow{p} \lambda_0. \tag{1.10}$$

##### 1.2.4 Confidence intervals for $\widehat{\lambda}_0$

We construct confidence intervals via subsampling with a Gaussian approximation. Draw $R$ subsamples $\{\mathcal{D}_r\}_{r=1}^R$ of size $b_n$ from the full sample, where $b_n \to \infty$ and $b_n/n \to 0$. For each subsample, compute the SIQReg estimate $\widehat{\lambda}_r$. Define

$$\bar{\lambda} = \frac{1}{R}\sum_{r=1}^R \widehat{\lambda}_r, \qquad \widehat{\sigma}^2 = \frac{1}{R-1}\sum_{r=1}^R (\widehat{\lambda}_r - \bar{\lambda})^2. \tag{1.11}$$

Then the $(1-\alpha)$-level confidence interval for $\lambda_0$ is

$$\left[\bar{\lambda} - z_{1-\alpha/2}\,\frac{\widehat{\sigma}}{\sqrt{R}},\ \ \bar{\lambda} + z_{1-\alpha/2}\,\frac{\widehat{\sigma}}{\sqrt{R}}\right], \tag{1.12}$$

where $z_{1-\alpha/2}$ is the $(1-\alpha/2)$-quantile of a standard normal random variable.

The justification proceeds as follows. By standard M-estimation theory (Van der Vaart, 2000), the full-sample estimator satisfies $\sqrt{n}(\widehat{\lambda} - \lambda_0) \xrightarrow{d} N(0, \sigma^2_{\lambda_0})$. By subsampling theory (Dimitris N. Politis, 1999), since $b_n/n \to 0$, each subsample estimator satisfies $\sqrt{b_n}(\widehat{\lambda}_r - \widehat{\lambda}) \xrightarrow{d} N(0, \sigma^2_{\lambda_0})$. Because $\widehat{\lambda} = \lambda_0 + O_p(n^{-1/2})$ and $b_n = o(n)$, we can replace the centering: $\sqrt{b_n}(\widehat{\lambda}_r - \lambda_0) \xrightarrow{d} N(0, \sigma^2_{\lambda_0})$. It follows that $\bar{\lambda} \sim N(\lambda_0, \sigma^2_{\lambda_0}/(b_n R))$ approximately, and $\widehat{\sigma}^2 \xrightarrow{p} \sigma^2_{\lambda_0}/b_n$. Therefore,

$$\frac{\sqrt{R}(\bar{\lambda} - \lambda_0)}{\widehat{\sigma}} \xrightarrow{d} N(0,1). \tag{1.13}$$

#### 1.3 Heterogeneity on the latent scale likely signals the genuine biology

After transforming the phenotype to the latent scale, detected quantile-dependent genetic effects are more likely to reflect biologically meaningful heterogeneity rather than measurement scale artifacts. We formalize this claim and characterize three biological mechanisms that can produce such heterogeneity.

Under (1.5), the genetic dependence of the conditional quantile comes from two parts: (1) the linear shift: $G\beta^*$; (2) any genetic dependence in the conditional quantile of error: $Q_{\varepsilon|G}(\tau)$. If the random error is independent of the genetic factor so that $Q_{\varepsilon|G}(\tau) = Q_{\varepsilon}(\tau)$, then

$$Q_{y_i^*|G_i,C_i}(\tau) = G_i\beta^* + C_i^\top \alpha^* + Q_{\varepsilon_i}(\tau), \tag{1.14}$$

so the marginal genetic effect is $\beta^*$ for each quantile level $\tau$. Therefore, detecting heterogeneous genetic effects on the SIQReg scale implies that $Q_{\varepsilon|G}(\tau)$ depends on $G$: the residual distribution varies across genotypes, a property that cannot be removed by any monotone rescaling. We now characterize three biological sources of this dependence.

**Gene-by-environment interaction (GxE).** If an unmeasured environment $E_i$ modifies the genetic effect:

$$\varepsilon_i = G_i E_i \gamma + \eta_i, \tag{1.15}$$

with i.i.d. $E_i \sim \mathcal{N}(0, \sigma_E^2)$ and $\eta_i \sim \mathcal{N}(0,1)$. This means the same variant has different effects across environmental exposures. Then $y|(G = g, C = c) \sim \mathcal{N}(g\beta^* + c^\top \alpha^*, \gamma^2 g^2 \sigma_E^2 + 1)$, so the conditional quantile of $y$ is:

$$Q_{y|G=g,C=c}(\tau) = g\beta^* + c^\top \alpha^* + z_\tau \sqrt{\gamma^2 g^2 \sigma_E^2 + 1}, \tag{1.16}$$

where $z_\tau = \Phi^{-1}(\tau)$ denotes the $\tau$-th quantile of the standard normal distribution, and the local genetic effect is:

$$\frac{\partial}{\partial g} Q_{y|G=g,C=c}(\tau) = \beta + z_\tau \frac{\gamma^2 g \sigma_E^2}{\sqrt{\gamma^2 g^2 \sigma_E^2 + 1}} := \beta(\tau). \tag{1.17}$$

If $\gamma = 0$, the genetic effect is quantile-independent. Otherwise, GxE induces quantile-dependent effects that grow in magnitude toward both tails.

**Genetic variance heterogeneity (vQTL).** If the variant directly affects phenotypic dispersion:

$$\varepsilon_i = \sigma(G_i)\, \eta_i, \tag{1.18}$$

with $\eta_i \sim \mathcal{N}(0, 1)$ and genotype-dependent scale $\sigma(g) > 0$. This model captures settings where a genetic variant influences not the mean phenotype but its variability. Under this model, $y^* \mid (G = g, C = c) \sim \mathcal{N}(g\beta^* + c^\top \alpha^*, \sigma^2(g))$, so the conditional quantile is

$$Q_{y^*|G=g,C=c}(\tau) = g\beta^* + c^\top \alpha^* + z_\tau \sigma(g), \tag{1.19}$$

and the local genetic effect is

$$\frac{\partial}{\partial g} Q_{y^*|G=g,C=c}(\tau) = \beta^* + z_\tau \frac{\partial \sigma(g)}{\partial g}. \tag{1.20}$$

If $\sigma'(g) = 0$, the genetic effect is quantile-independent. Otherwise, variance heterogeneity produces effects that are not constant across quantile levels.

**Gene-by-gene interaction (epistasis).** If a second locus $H_i$ interacts with $G_i$:

$$\varepsilon_i = H_i(\delta + G_i \phi) + \eta_i, \tag{1.21}$$

with i.i.d. $H_i \sim \mathcal{N}(0, \sigma_H^2)$ and $\eta_i \sim \mathcal{N}(0, 1)$. This means the effect of $G_i$ depends on the genetic background at $H_i$. Under this model, $y^* \mid (G = g, C = c) \sim \mathcal{N}(g\beta^* + c^\top \alpha^*, (\delta + g\phi)^2 \sigma_H^2 + \mathbf{1})$, so the conditional quantile is

$$Q_{y^*|G=g,C=c}(\tau) = g\beta^* + c^\top \alpha^* + z_\tau \sqrt{(\delta + g\phi)^2 \sigma_H^2 + 1}, \tag{1.22}$$

and the local genetic effect is

$$\frac{\partial}{\partial g} Q_{y^*|G=g,C=c}(\tau) = \beta^* + z_\tau \frac{\phi(\delta + g\phi)\sigma_H^2}{\sqrt{(\delta + g\phi)^2 \sigma_H^2 + 1}}. \tag{1.23}$$

If $\phi = 0$, the genetic effect is quantile-independent. Otherwise, epistasis induces quantile-dependent effects whose pattern depends on the sign and magnitude of $\delta + g\phi$.

##### 1.4 Latent scale transformation improves power

We now investigate the impact of transforming the phenotype to the latent scale on the additive association test. We consider two testing strategies:

- **Latent scale**: Regress $y_i^*$ on $G_i$.
- **Default scale**: Regress $y_i$ on $G_i$ directly.

We will show that the latent-scale test has uniformly higher power. For the sake of clarity, we consider the latent phenotype model without covariates and normal errors:

$$y_i^* = G_i \beta^* + e_i, \quad e_i \sim \mathcal{N}(0, \sigma_e^2), \tag{1.24}$$

where $E[G_i] = 0$ and $G_i \perp\!\!\!\perp e_i$. The observed phenotype $y_i$ is related to the latent phenotype through a general nonlinear transformation:

$$y_i = f_{\lambda_0}^{-1}(y_i^*). \tag{1.25}$$

We first consider the association test on the latent scale. Let $A = \mathrm{E}[G_i^2]$, standard asymptotic theory gives that:

$$\sqrt{n}(\widehat{\beta}^* - \beta^*) \xrightarrow{d} \mathcal{N}\left(0,\ \frac{\sigma_e^2}{A}\right) := \mathcal{N}(0, V_{\text{trans}}). \tag{1.26}$$

For testing $H_0 : \beta^* = 0$ under local alternatives $H_1 : \beta^* = h/\sqrt{n}$, the $\alpha$ level power function of the association test on the latent scale is:

$$\pi_{\text{latent}}(h) = 1 - \Phi\left(z_{1-\alpha} - \frac{h}{\sqrt{V_{\text{trans}}}}\right) = 1 - \Phi\left(z_{1-\alpha} - h\sqrt{\frac{A}{\sigma_e^2}}\right). \tag{1.27}$$

We now consider the association test on the default scale where the estimator is $\widehat{\beta}^{(\text{obs})} = \frac{\sum G_i y_i}{\sum G_i^2}$. Write:

$$\widehat{\beta}^{(\text{obs})} - \beta^{(\text{obs})} = \frac{\frac{1}{n}\sum G_i y_i}{\frac{1}{n}\sum G_i^2} - \frac{E[G_i y_i]}{E[G_i^2]} = \frac{E[G_i^2] \cdot \frac{1}{n}\sum G_i y_i - E[G_i y_i] \cdot \frac{1}{n}\sum G_i^2}{E[G_i^2] \cdot \frac{1}{n}\sum G_i^2}.$$

Multiplying by $\sqrt{n}$ and applying Slutsky's theorem, the asymptotic distribution is determined by the numerator:

$$\frac{1}{\sqrt{n}}\sum_{i=1}^{n}\big(A \cdot G_i y_i - E[G_i y_i] \cdot G_i^2\big) \xrightarrow{d} \mathcal{N}\big(0,\ \text{Var}(A \cdot G_i y_i - E[G_i y_i] \cdot G_i^2)\big).$$

The limiting variance expands as:

$$\text{Var}(A \cdot G_i y_i - E[G_i y_i] \cdot G_i^2) = A^2\,\text{Var}(G_i y_i) + (E[G_i y_i])^2\,\text{Var}(G_i^2) - 2A\,E[G_i y_i]\,\text{Cov}(G_i y_i,\, G_i^2).$$

For $\text{Var}(G_i y_i)$, since $y_i = f_{\lambda_0}^{-1}(G_i\beta^* + e_i)$, expanding $y_i^2$ in the small quantity $G_i\beta^*$:

$$y_i^2 = (f_{\lambda_0}^{-1}(e_i))^2 + 2G_i\beta^* \cdot f_{\lambda_0}^{-1}(e_i)(f_{\lambda_0}^{-1})'(e_i) + O(G_i^2\beta^{*2}),$$

so that

$$E[G_i^2 y_i^2] = E[G_i^2] \cdot E[(f_{\lambda_0}^{-1}(e_i))^2] + O(\beta^*) = A \cdot E[(f_{\lambda_0}^{-1}(e_i))^2] + O(\beta^*).$$

For $E[G_i y_i]$, expansion gives:

$$y_i = f_{\lambda_0}^{-1}(G_i\beta^* + e_i) = f_{\lambda_0}^{-1}(e_i) + G_i\beta^* \cdot (f_{\lambda_0}^{-1})'(e_i) + O(G_i^2\beta^{*2}).$$

Multiplying by $G_i$ and taking expectations gives:

$$E[G_i y_i] = \beta^* \cdot E[G_i^2] \cdot E[(f_{\lambda_0}^{-1})'(e_i)] + O(\beta^{*2}) = A \cdot c_{\lambda_0} \cdot \beta^* + O(\beta^{*2}),$$

where $c_{\lambda_0} := E[(f_{\lambda_0}^{-1})'(e_i)]$. Under local alternatives $\beta^* = h/\sqrt{n}$, we have $E[G_i y_i] = Ac_{\lambda_0}\beta^* = O(n^{-1/2})$. Therefore, terms involving $(E[G_i y_i])^2 = O(n^{-1})$ and $E[G_i y_i] = O(n^{-1/2})$ are negligible, so that:

$$\mathrm{Var}(G_i y_i) = E[G_i^2 y_i^2] - (E[G_i y_i])^2 \to A \cdot E[(f_{\lambda_0}^{-1}(e))^2].$$

Therefore,

$$\mathrm{Var}(A \cdot G_i y_i - E[G_i y_i] \cdot G_i^2) \to A^3 \cdot E[(h_{\lambda_0}^{-1}(e))^2].$$

The asymptotic distribution of $\widehat{\beta}^{(\mathrm{obs})}$ is therefore:

$$\sqrt{n}\big(\widehat{\beta}^{(\mathrm{obs})} - \beta^{(\mathrm{obs})}\big) \xrightarrow{d} \mathcal{N}\left(0,\ \frac{E[(f_{\lambda_0}^{-1}(e))^2]}{A}\right) := \mathcal{N}(0, V_{\mathrm{obs}}). \tag{1.28}$$

Finally, the power function of the association test on the default scale is:

$$\pi_{\mathrm{orig}}(h) = 1 - \Phi\left(z_{1-\alpha} - \frac{c_{\lambda_0} h}{\sqrt{V_{\mathrm{obs}}}}\right) = 1 - \Phi\left(z_{1-\alpha} - c_{\lambda_0} h \sqrt{\frac{A}{E[(f_{\lambda_0}^{-1}(e))^2]}}\right). \tag{1.29}$$

Now we compare the power functions of two approaches in the following theorem:

**Theorem 1.1.** *For any fixed $h > 0$,*

$$\pi_{\mathrm{trans}}(h) \geq \pi_{\mathrm{obs}}(h),$$

*with equality if and only if $\lambda_0 = 1$.*

*Proof.* Since $e_i \sim \mathcal{N}(0, \sigma_e^2)$, Stein's lemma gives:

$$E[(f_{\lambda_0}^{-1})'(e)] = \frac{E[e \cdot f_{\lambda_0}^{-1}(e)]}{\sigma_e^2} = \frac{\mathrm{Cov}(e,\ f_{\lambda_0}^{-1}(e))}{\sigma_e^2},$$

so that

$$c_{\lambda_0} \sigma_e^2 = \mathrm{Cov}(e,\ f_{\lambda_0}^{-1}(e)). \tag{1.30}$$

Applying the Cauchy–Schwarz inequality to (1.30) gives

$$c_{\lambda_0}^2 \sigma_e^4 = \big[\mathrm{Cov}(e,\ h_{\lambda_0}^{-1}(e))\big]^2 \leq \mathrm{Var}(e) \cdot \mathrm{Var}(h_{\lambda_0}^{-1}(e)) = \sigma_e^2 \cdot \mathrm{Var}(h_{\lambda_0}^{-1}(e)).$$

Dividing both sides by $\sigma_e^2$:

$$c_{\lambda_0}^2 \sigma_e^2 \leq \mathrm{Var}(h_{\lambda_0}^{-1}(e)) \leq E[(h_{\lambda_0}^{-1}(e))^2].$$

This gives

$$\frac{c_{\lambda_0}^2 \sigma_e^2}{E[(h_{\lambda_0}^{-1}(e))^2]} \le 1, \tag{1.31}$$

Since $\Phi$ is monotonically increasing, $\pi_{\text{trans}}(h) \ge \pi_{\text{obs}}(h)$ if and only if

$$\frac{h}{\sqrt{V_{\text{trans}}}} \ge \frac{c_{\lambda_0} h}{\sqrt{V_{\text{obs}}}} \quad \iff \quad \frac{c_{\lambda_0}^2 \sigma_e^2}{E[(f_{\lambda_0}^{-1}(e))^2]} \le 1,$$

which is verified by (1.31). □

#### 1.5 A mathematical model explaining pervasive concave latent transformations

SIQReg identifies $\lambda_0 < 1$ for 24/25 UK Biobank traits examined. We propose a generative model that explains this finding by positing that genetic variants affect phenotypes through a mixture of additive and multiplicative channels.

Suppose the phenotype on the default scale is generated as

$$y = \prod_{j=1}^{M} (1 + a_j g_j)^{1-\theta_j} e^{\theta_j b_j g_j}, \tag{1.32}$$

where $g_j$ is the $j$-th variant, $a_j$ is the additive effect, $b_j$ is the multiplicative effect, and $\theta_j \in [0, 1]$ is a continuous index of additivity versus multiplicativity:

- When $\theta_j = 0$ for each $j$, $y = \prod_{j=1}^{M}(1 + a_j g_j) \approx 1 + \sum_{j=1}^{M} a_j g_j$ so that all variants affect traits additively.
- When $\theta_j = 1$ for each $j$, $y = e^{\sum_{j=1}^{M} b_j g_j}$ so that all variants affect traits multiplicatively.

We now show that this model can naturally yield $\lambda_0 < 1$. First, note that

$$\begin{aligned}
\log y &= \sum_{j=1}^{M} (1 - \theta_j) \log(1 + a_j g_j) + \sum_{j=1}^{M} \theta_j b_j g_j \\
&\approx \sum_{j=1}^{M} (1 - \theta_j) \left( a_j g_j - \frac{1}{2} a_j^2 g_j^2 \right) + \sum_{j=1}^{M} \theta_j b_j g_j \\
&= \sum_{j=1}^{M} \left[ (1 - \theta_j) a_j + \theta_j b_j \right] g_j - \frac{1}{2} \sum_{j=1}^{M} (1 - \theta_j) a_j^2 g_j^2 \\
&:= I_1 - \frac{1}{2} I_2,
\end{aligned}$$

where the first approximation follows from Taylor expansions. As a result,

$$y^*(\lambda) = \frac{y^\lambda - 1}{\lambda} = \frac{e^{\lambda \log y} - 1}{\lambda} \approx \log y + \frac{\lambda}{2}\log^2 y \approx I_1 - \frac{1}{2}I_2 + \frac{\lambda}{2}\left(I_1 - \frac{1}{2}I_2\right)^2 \approx I_1 - \frac{1}{2}I_2 + \frac{\lambda}{2}I_1^2$$

assuming cross-locus interactions are negligible, so that

$$y^*(\lambda) \approx \sum_{j=1}^{M} w_j g_j + \frac{1}{2}\sum_{j=1}^{M}[\lambda w_j^2 - (1-\theta_j)a_j^2]g_j^2 \tag{1.33}$$

where $w_j := (1-\theta_j)a_j + \theta_j b_j$. Therefore, the optimal $\lambda$ that makes latent phenotype additive satisfies:

$$\begin{aligned}\lambda_0 &= \arg\min_{\lambda} \sum_{j=1}^{M} s_j[\lambda w_j^2 - (1-\theta_j)a_j^2]^2 \quad \text{where } s_j \propto \mathrm{Var}(g_j^2) \\ &= \frac{\sum_{j=1}^{M} s_j w_j^2 (1-\theta_j)a_j^2}{\sum_{j=1}^{M} s_j w_j^4}.\end{aligned}$$

We further assume that the additive effect and multiplicative effect of each variant is the same, i.e., $a_j = b_j$, then

$$\lambda_0 = 1 - \frac{\sum_{j=1}^{M} s_j \theta_j a_j^4}{\sum_{j=1}^{M} s_j a_j^4} \in [0,1), \tag{1.34}$$

and $\lambda_0 = 0$ iff $\theta_j = 1$ for all $j$. More generally, $a_j = b_j$ is not required but made for analytical clarity; (1.34) holds when $w_j^2 > (1-\theta_j)a_j^2$ on average. This result says that $\lambda_0$ is one minus the weighted average multiplicative index across the genome. Several features of this result are notable:

First, $\lambda_0$ is not a property of individual variants but an aggregate over the full polygenicity of a trait. A trait with $\lambda_0 = 0.5$ is not "half additive and half multiplicative" at each locus but the average channel usage is balanced between the two modes across the full set of contributing variants. Individual variants may be predominantly additive or predominantly multiplicative, but the aggregate determines $\lambda_0$.

Second, the model provides a possible answer to the longstanding additive-versus-multiplicative debate in quantitative genetics. Under our model, the question is not which model is correct, but what mixture of the two best describes each trait. The latent scale parameter $\lambda_0$ provides a continuous, empirically estimable answer.

#### 2 Supplementary Figures

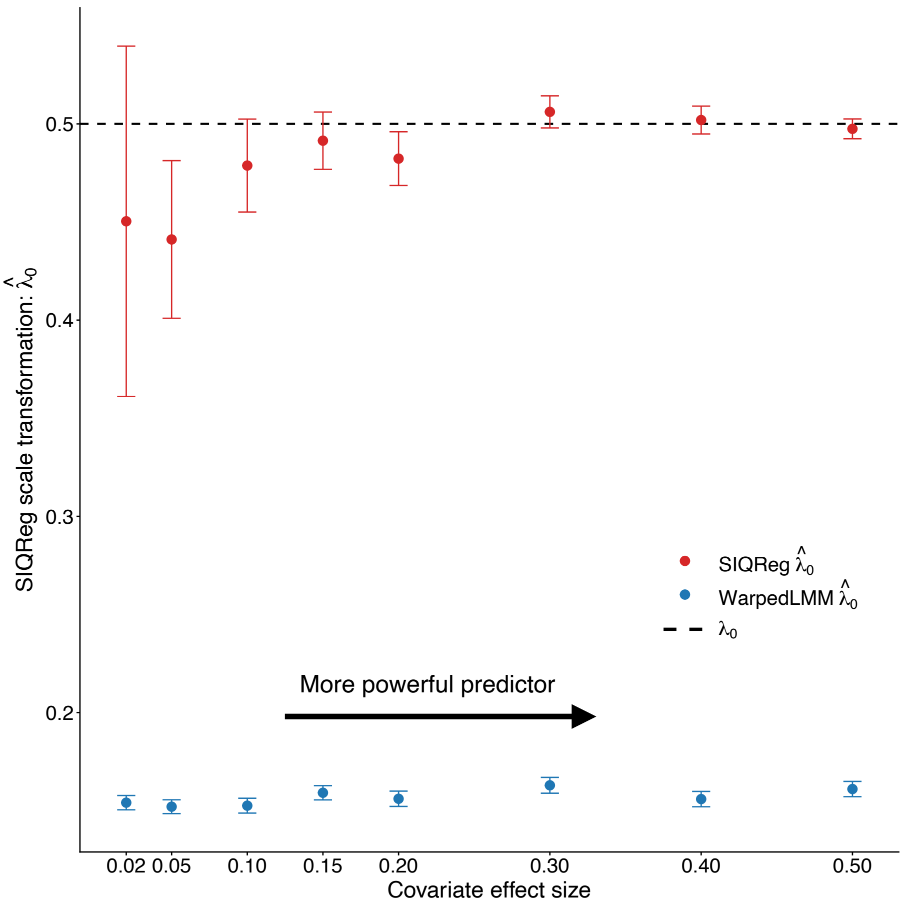

Supplementary Figure 1: Impact of predictor strength on SIQReg estimates $\widehat{\lambda}_0$

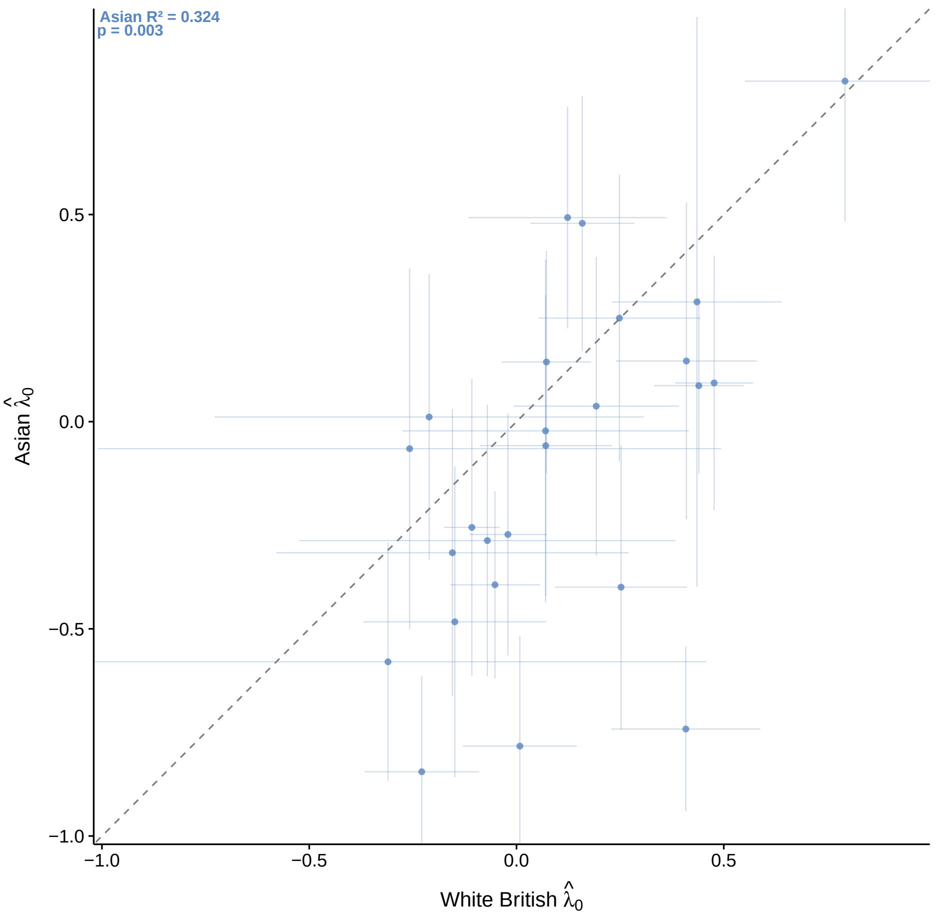

Supplementary Figure 2: SIQReg scale transformation in White British versus in Asian

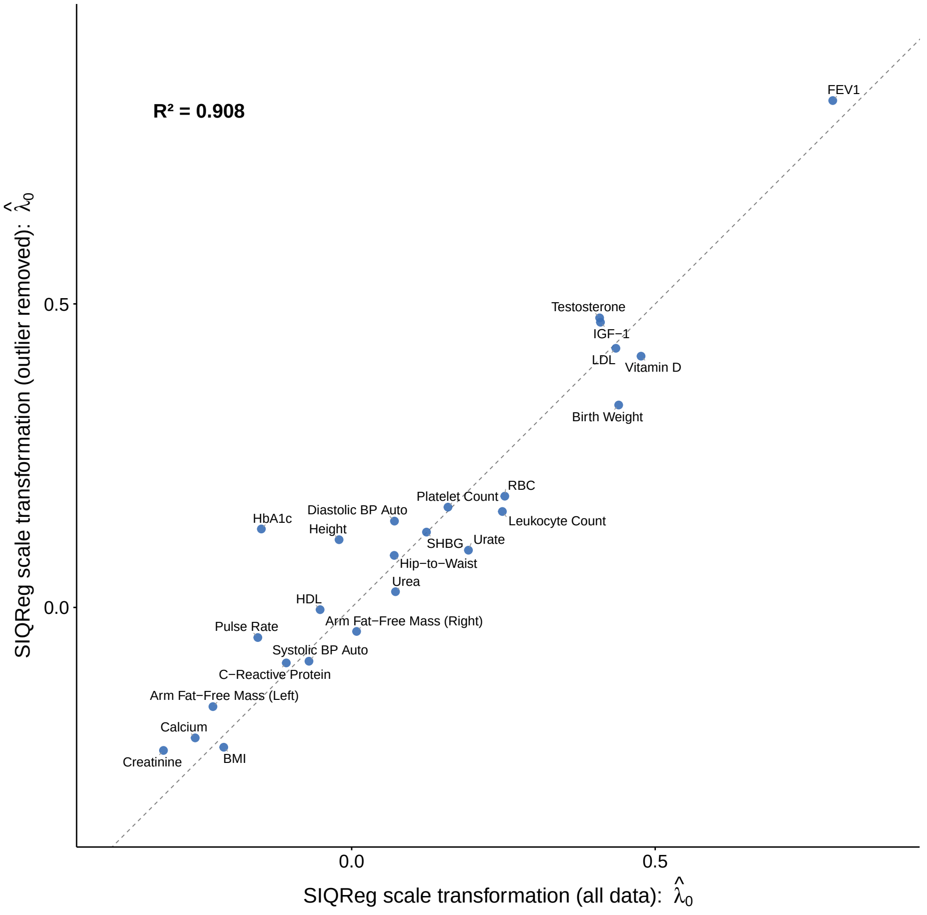

Supplementary Figure 3: SIQReg scale transformation with 6 s.d. phenotypic values excluded

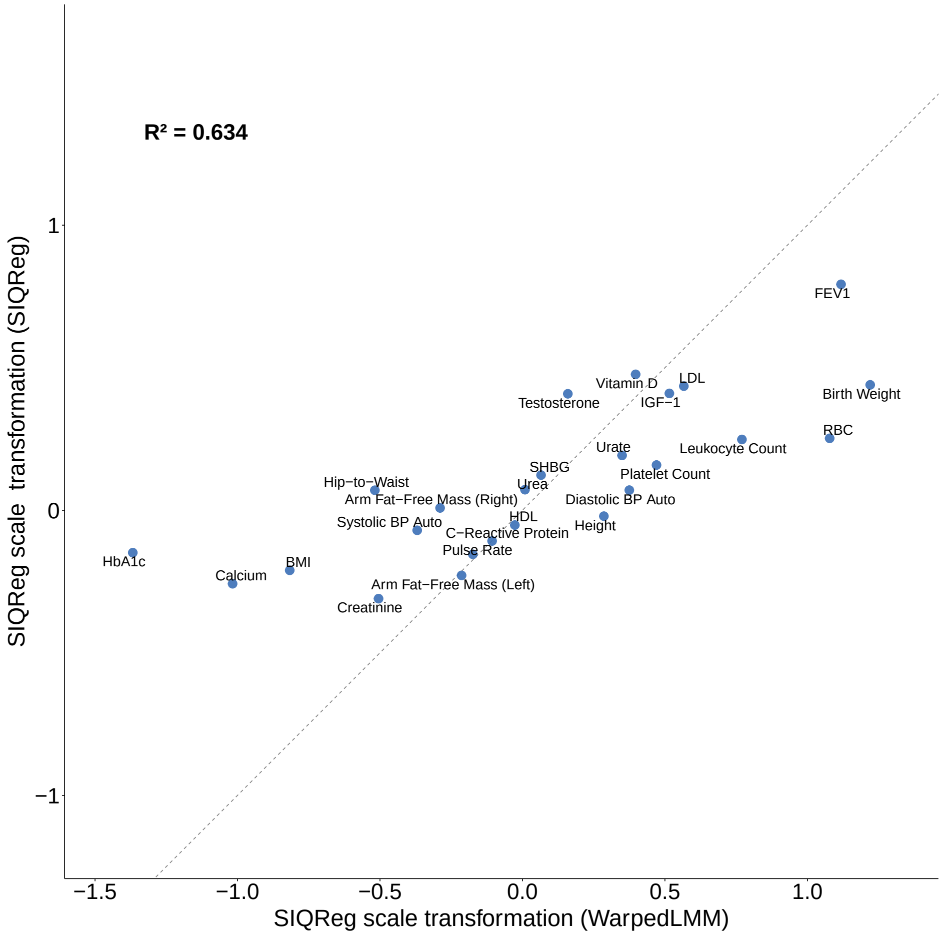

Supplementary Figure 4: SIQReg scale transformation versus WarpedLMM scale transformation

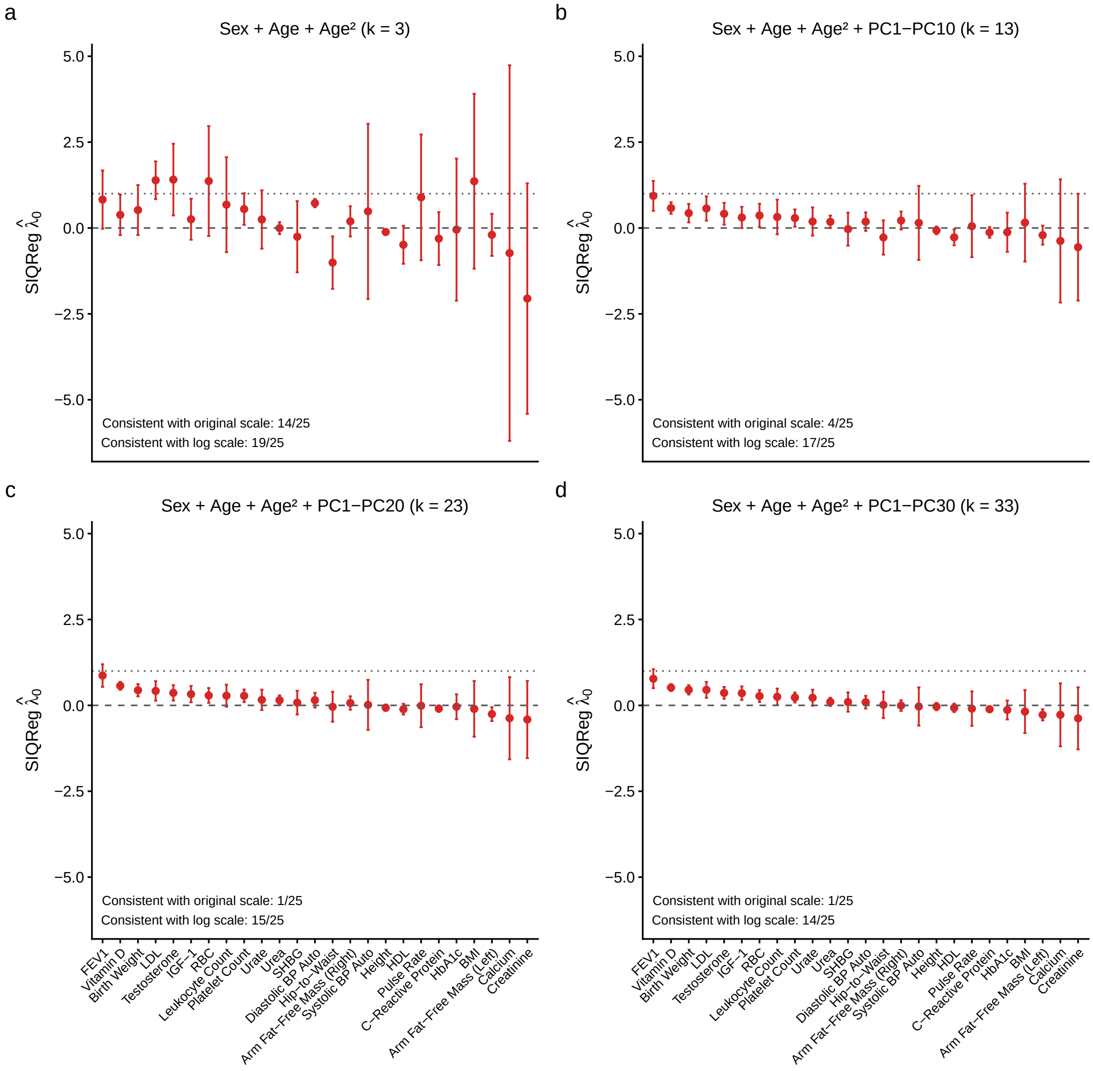

Supplementary Figure 5: SIQReg scale transformation with different number of covariates

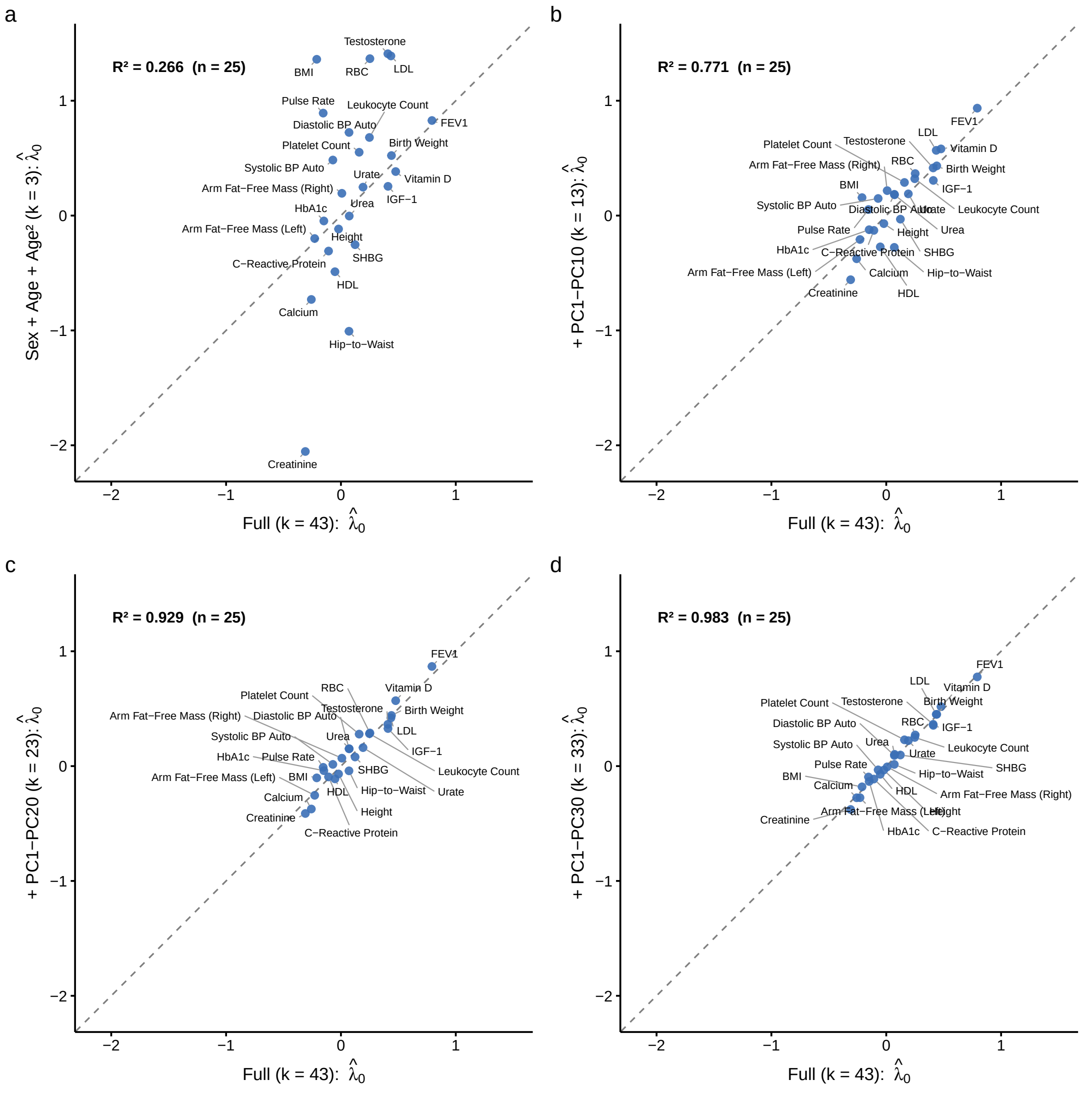

Supplementary Figure 6: SIQReg scale transformation concordance under different number of covariates

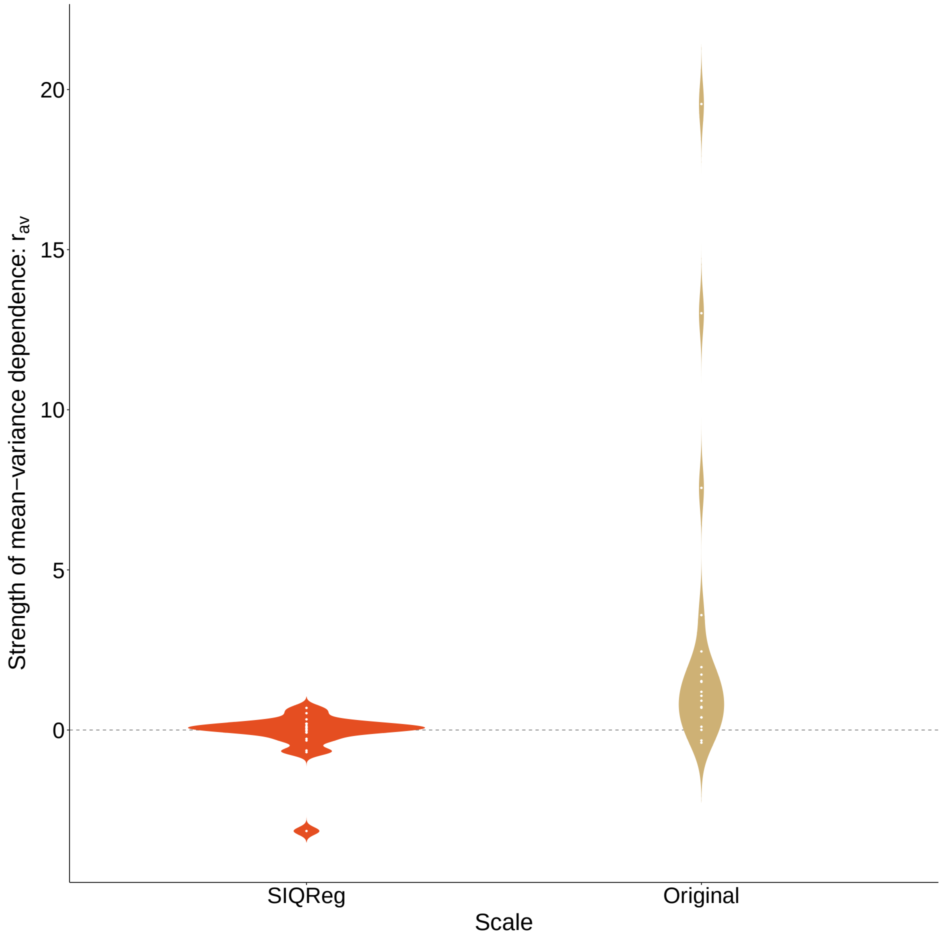

Supplementary Figure 7: Mean-variance dependence $r_{\mathrm{av}}$ on the default and SIQReg scale

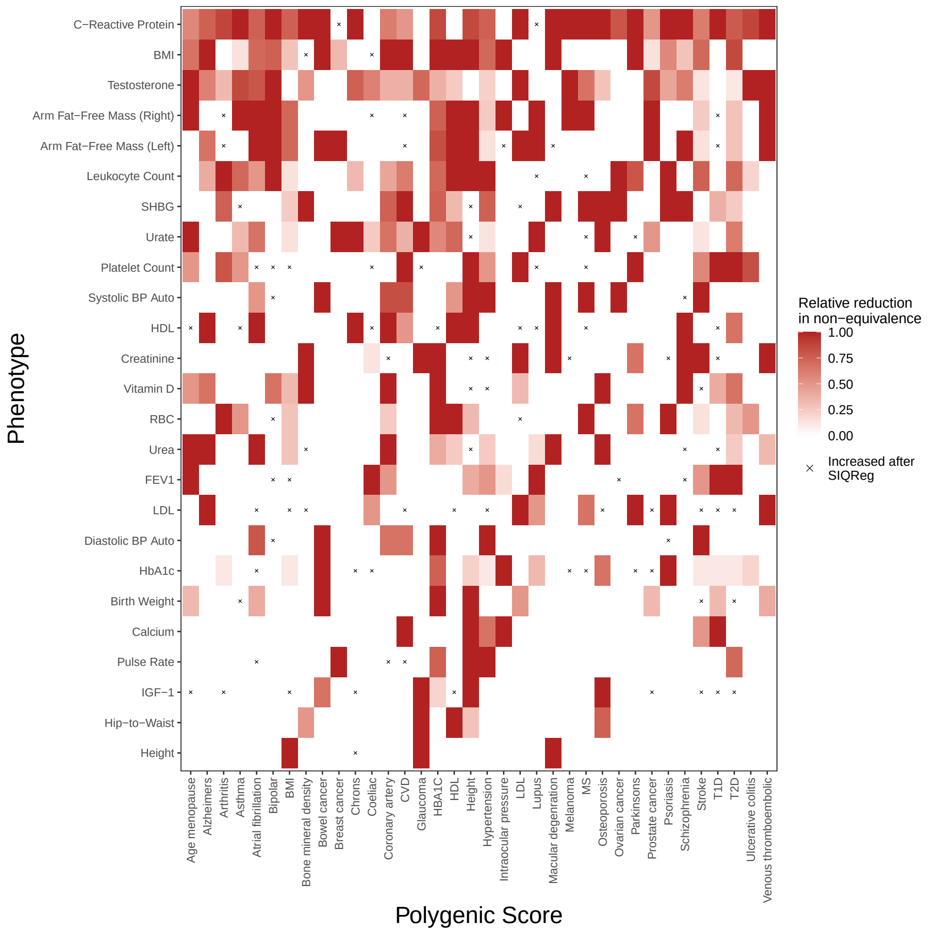

Supplementary Figure 8: Heatmap of nonequivalent PGS changes after SIQReg transformation

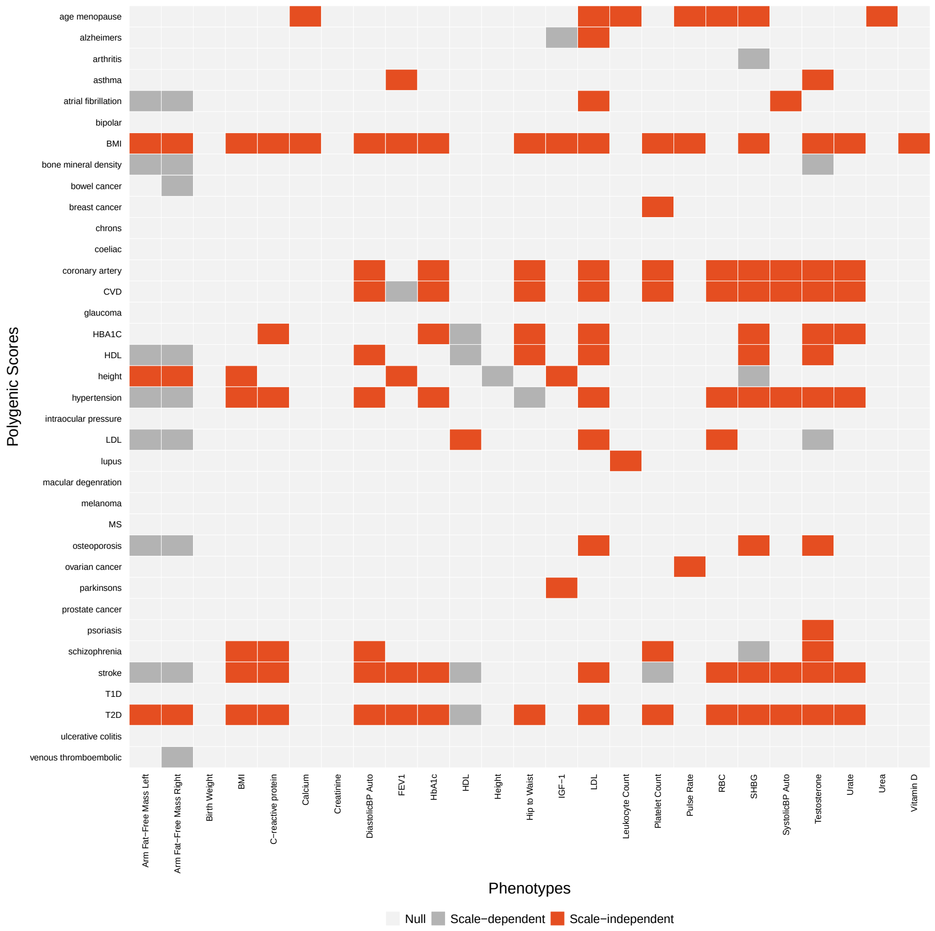

Supplementary Figure 9: Heatmap of PGSxSex across all 25 complex traits

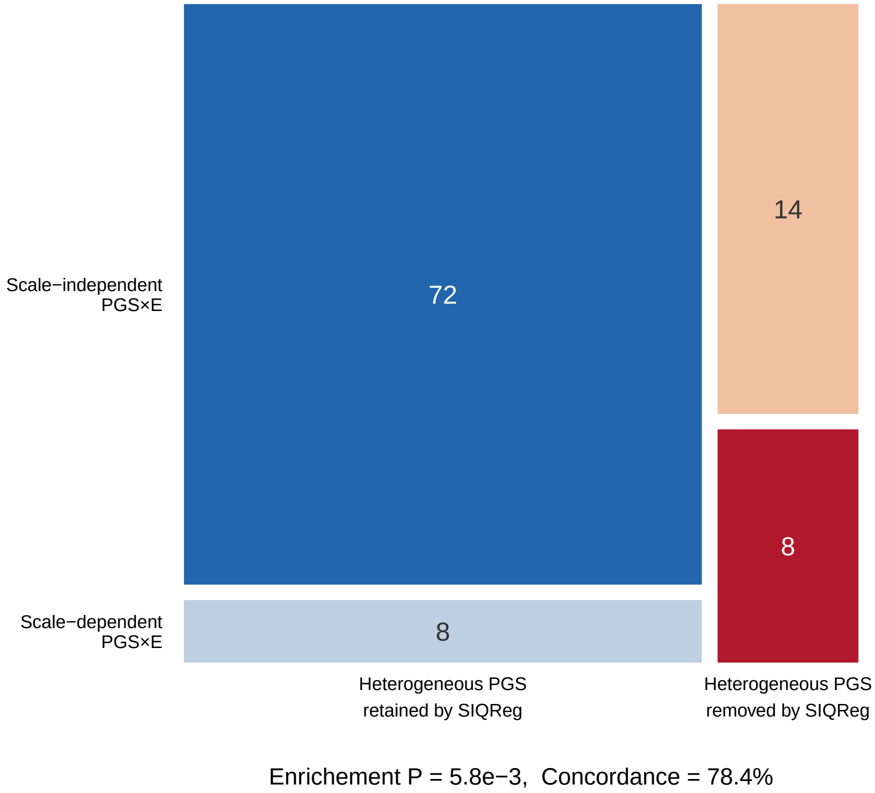

Supplementary Figure 10: Number of PGSxSex interactions versus number of heterogeneous PGS across all significant PGS-trait pairs

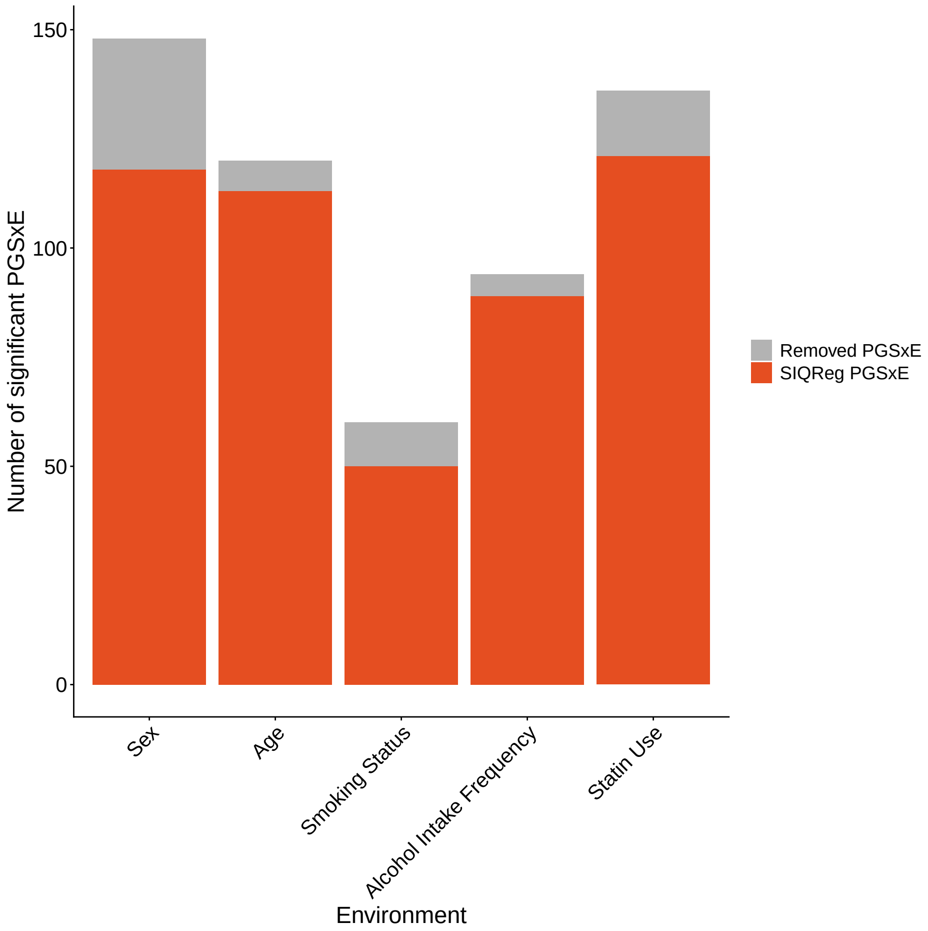

Supplementary Figure 11: Number of PGSxE interactions found across PGS-phenotypes pairs for each environment

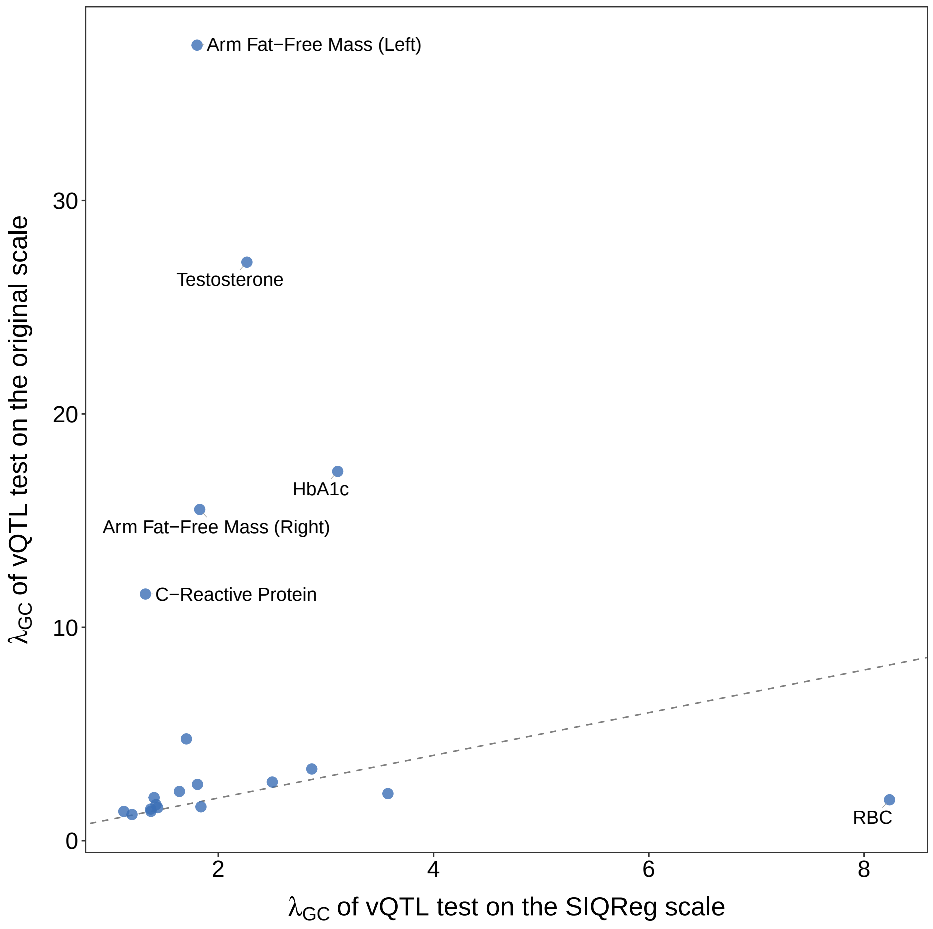

Supplementary Figure 12: vQTL test $\lambda_{GC}$ on the default and SIQReg scale

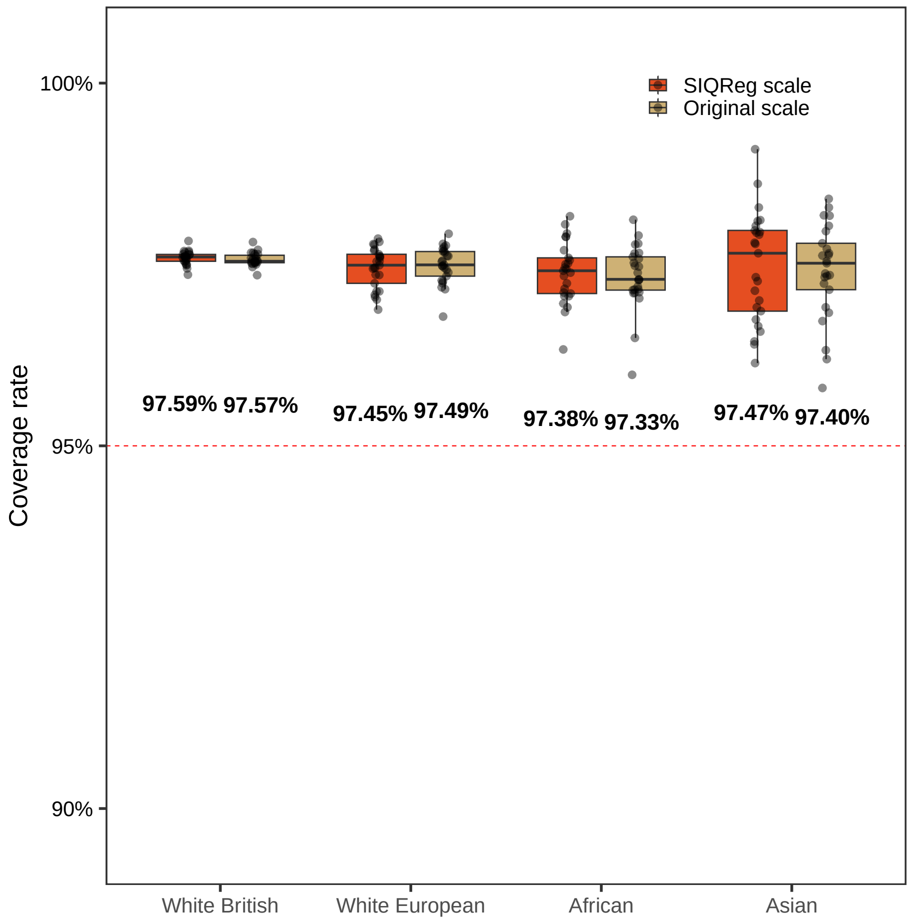

Supplementary Figure 13: PredInterval coverage rate across phenotypes on the default and SIQReg scale in four ancestry groups

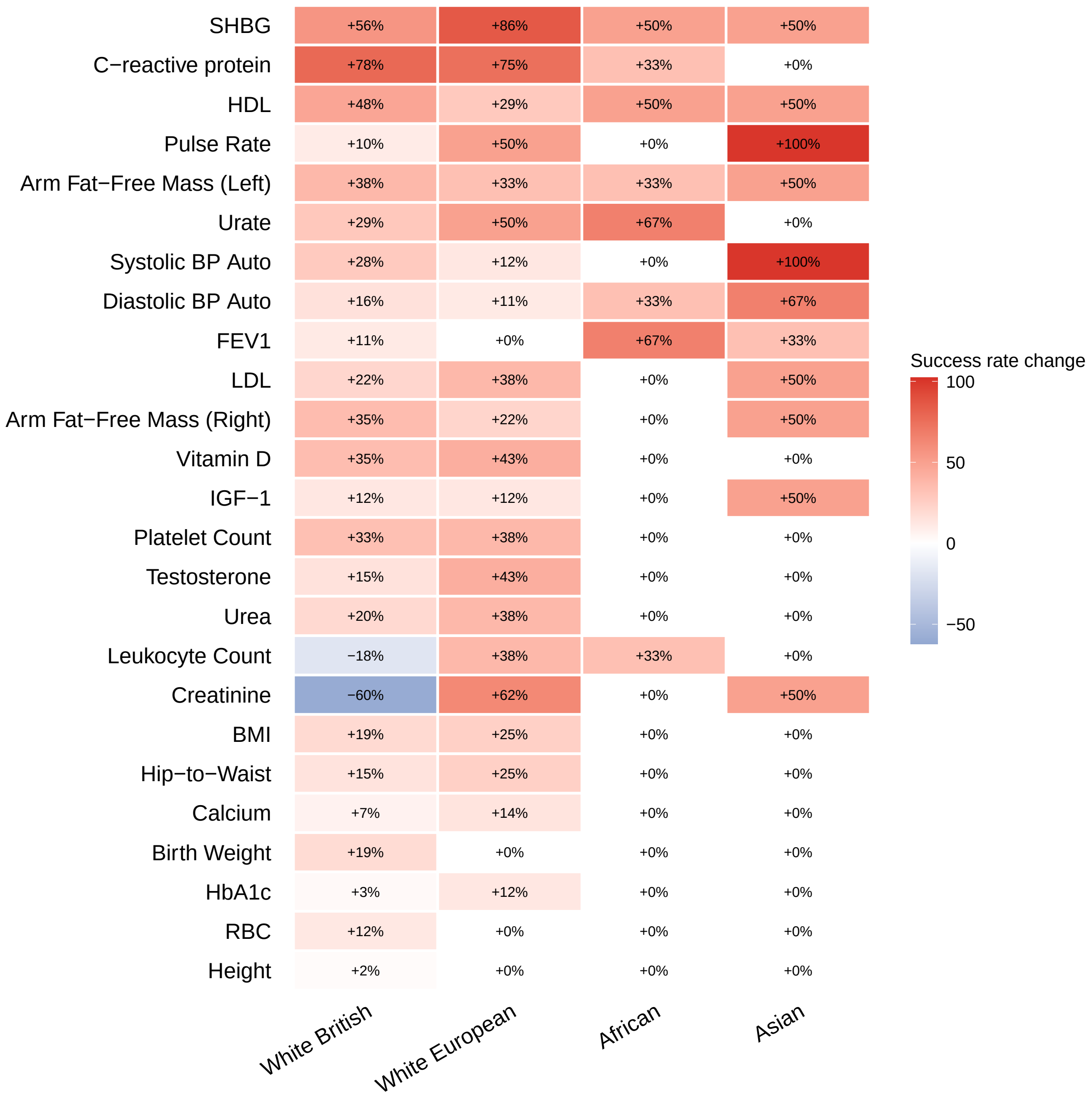

Supplementary Figure 14: PredInterval success rate change for identifying high-risk individuals (top 0.1%) in four ancestry groups

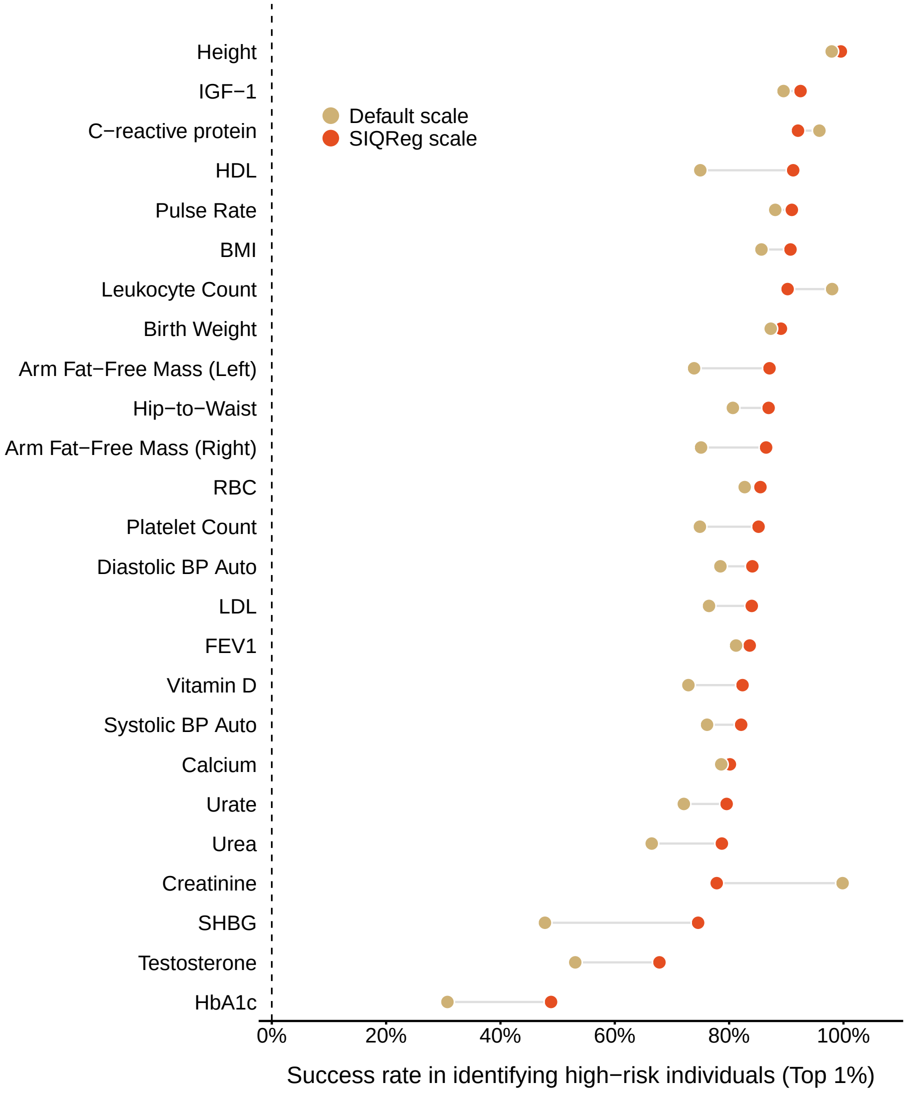

Supplementary Figure 15: Success rate for identifying high-risk individuals (top 1%) using PGS on the default and SIQReg scales

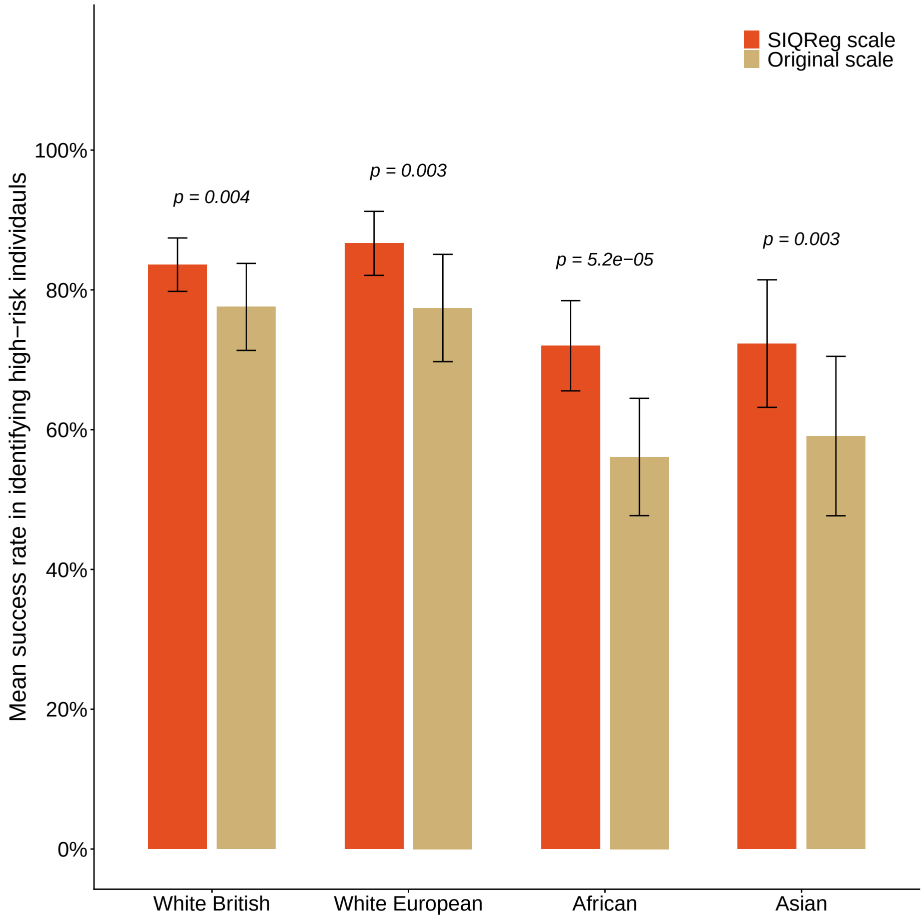

Supplementary Figure 16: Mean success rate for high-risk identification (top 1%) across traits in four ancestry groups

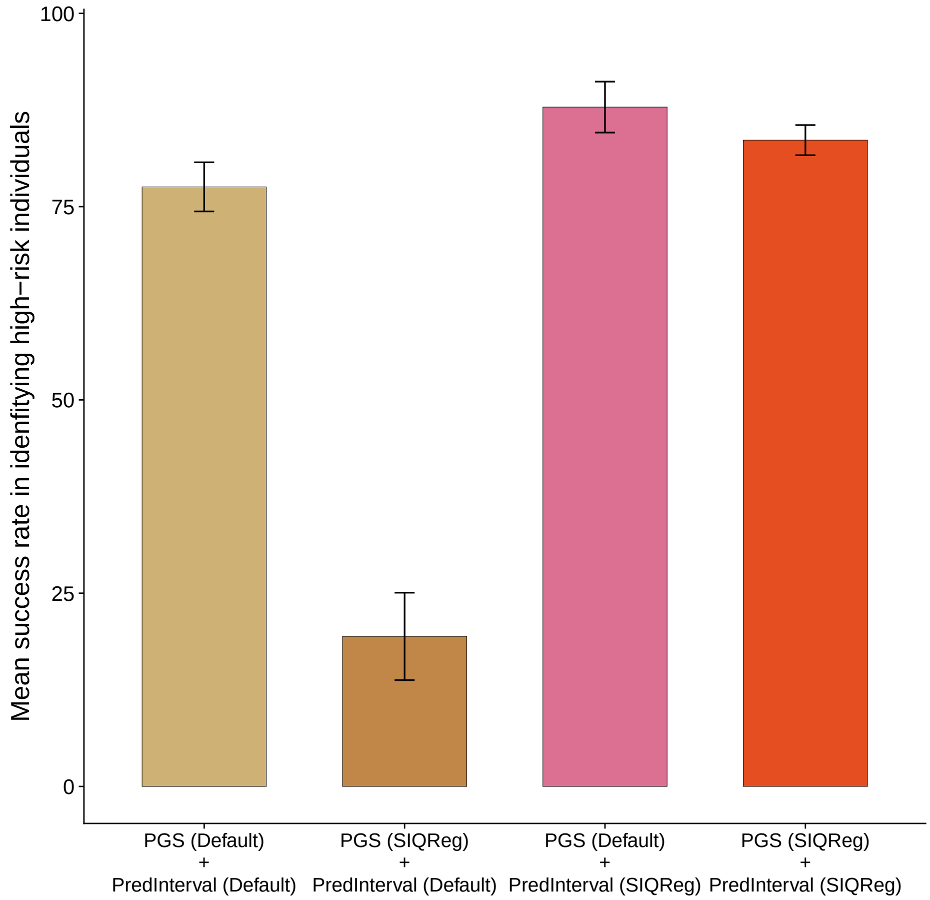

Supplementary Figure 17: Mean success rate across 25 traits for identifying high-risk individuals (top 1%) under four combinations of phenotype scale used for PGS construction and prediction interval estimation: Default PGS + Default prediction interval, SIQReg PGS + Default prediction interval, Default PGS + SIQReg prediction interval, and SIQReg PGS + SIQReg prediction interval.

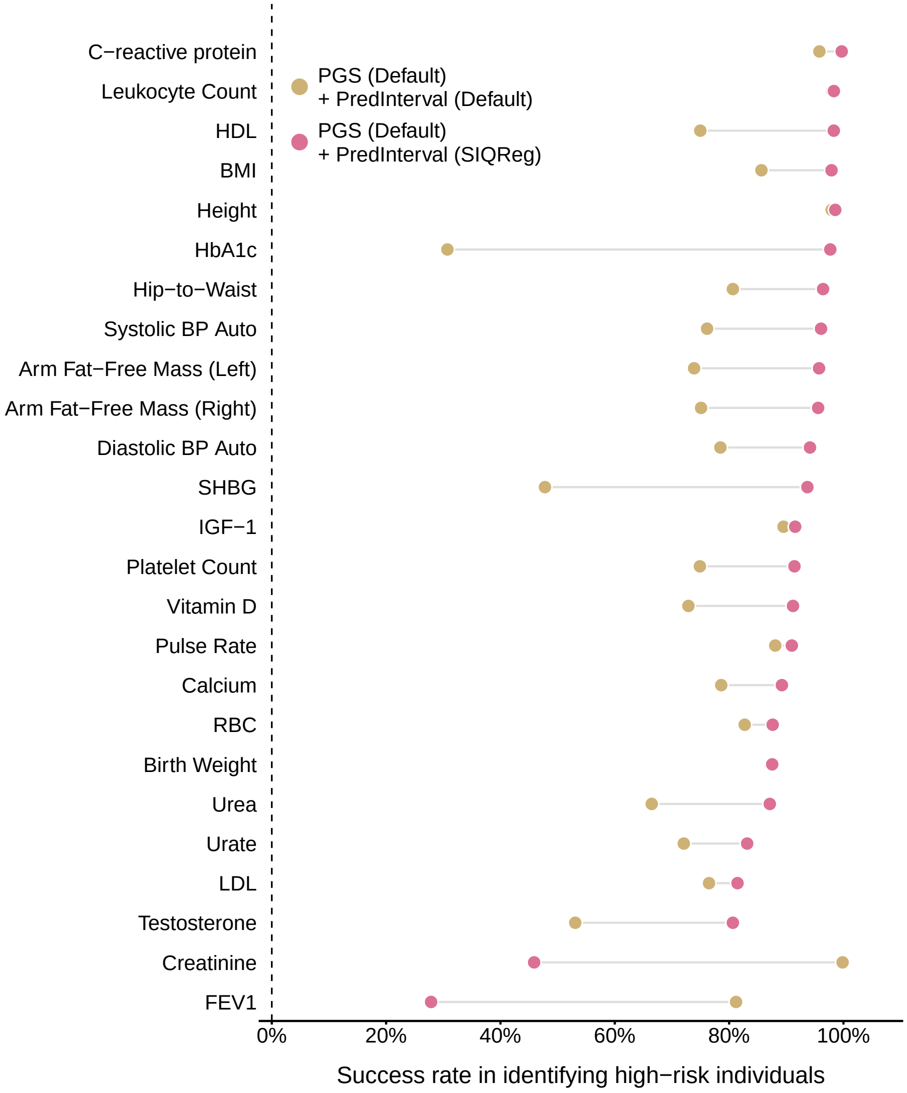

Supplementary Figure 18: Trait-by-trait comparison between Default PGS + Default prediction interval and Default PGS + SIQReg prediction interval (top 1%). Holding the PGS fixed on the default scale shows that part of the improvement in tail identification is attributable specifically to better-calibrated prediction intervals on the SIQReg scale.
